# The α4 nicotinic acetylcholine receptor is necessary for the initiation of organophosphate-induced neuronal hyperexcitability

**DOI:** 10.1101/2024.01.24.576980

**Authors:** Peter M. Andrew, Wei Feng, Jonas J. Calsbeek, Shane P. Antrobus, Gennady A. Cherednychenko, Jeremy A. MacMahon JA, Pedro N. Bernardino, Xiuzhen Liu, Danielle J. Harvey, Pamela J. Lein, Isaac N. Pessah

## Abstract

Acute intoxication with organophosphorus (OP) cholinesterase inhibitors can produce seizures that rapidly progress to life-threatening *status epilepticus*. Significant research effort has been invested investigating the involvement of muscarinic acetylcholine receptors (mAChRs) in OP-induced seizure activity. In contrast, there has been far less effort focused on nicotinic AChRs (nAChRs) in this context. Here, we address this data gap using a combination of *in vitro* and *in vivo* models. Pharmacological antagonism and genetic deletion of α4, but not α7, nAChR subunits prevented or significantly attenuated OP-induced electrical spike activity in acute hippocampal slices and seizure activity in mice, indicating that α4 nAChR activation is necessary for neuronal hyperexcitability triggered by acute OP exposures. These findings not only suggest that therapeutic strategies for inhibiting the α4 nAChR subunit warrant further investigation as prophylactic and acute treatments for acute OP-induced seizures, but also provide mechanistic insight into the role of the nicotinic cholinergic system in seizure generation.

## Introduction

Organophosphorus cholinesterase inhibitors (OPs) are a class of toxic chemicals widely used as insecticides that have also been weaponized for use against military and civilian targets (Costa, 2018; Jett and Spriggs, 2020). OPs pose a significant risk to public health and national security (Jett and Spriggs, 2020; Steindl et al., 2021; Vale et al., 2018). OP insecticides account for over 100,000 annual deaths globally, primarily as the result of accidental exposures or suicide attempts (Jett et al., 2020; Mew et al., 2017). In addition, there have been numerous high-profile instances of malicious OP nerve agent use, including the 1995 Tokyo subway sarin gas attack, more recent civilian exposures in the Syrian Civil War, and targeted assassination attempts on North Korean and Russian dissidents (Jett and Spriggs, 2020).

The canonical mechanism of acute OP neurotoxicity is inhibition of acetylcholinesterase (AChE; EC 3.1.1.7), which results in hyperstimulation of muscarinic and nicotinic acetylcholine receptors (AChR) in the peripheral and central nervous systems (Tsai and Lein, 2021). Acute inhibition of AChE by ≥ 60-70% causes “cholinergic crisis”, a clinical toxidrome characterized by muscle fasciculations and weakness, parasympathomimetic signs, depression of respiratory control centers in the brainstem, and seizures that can progress to life-threatening *status epilepticus* (SE) (Pereira et al., 2014; Richardson et al., 2019). Current standard of care for cholinergic crisis includes atropine to inhibit muscarinic acetylcholine receptors (mAChR), oximes to reactivate AChE that has not yet aged, and benzodiazepines for seizure management (Newmark, 2019). While prompt therapeutic intervention improves survival following acute OP intoxication, survivors often develop significant long-term morbidity, including progressive brain damage, electroencephalographic abnormalities and cognitive impairment (Chen, 2012; Chuang et al., 2019; Dassanayake et al., 2007, 2008; de Araujo Furtado et al., 2012; Figueiredo et al., 2018; Jett et al., 2020; Loh et al., 2010; Newmark, 2019; Yamasue et al., 2007). The severity and extent of chronic neurotoxicity observed following acute OP intoxication is proportional to the duration and intensity of the acute OP-induced SE (Figueiredo et al., 2018; Hobson et al., 2017; Shih et al., 2003). Thus, a reasonable therapeutic strategy for mitigating adverse neurological sequelae following acute OP intoxication is to prevent the onset or minimize the duration and/or severity of OP-induced SE. However, identifying more effective therapeutic targets has been stymied by the lack of a comprehensive understanding of the mechanisms involved in seizure generation following acute OP intoxication.

Most mechanistic studies of OP-induced seizures have focused on the importance of mAChR signaling in seizure initiation and the rapid transition to excessive glutamatergic signaling (McDonough and Shih, 1997; Shih et al., 1991; Tsai and Lein, 2021) with subsequent downregulation of inhibitory GABA receptors (Goodkin et al., 2008; Wasterlain et al., 2009) to maintain seizures. Consequently, the development of therapeutic strategies for terminating OP-induced seizures has largely focused on pharmacological antagonism of muscarinic and glutamatergic signaling and positive allosteric modulation of GABAergic signaling. In contrast, the role of nicotinic AChR (nAChR) signaling has not been extensively investigated as a therapeutic target in OP-induced seizures. This may be due in part to the fact that appropriate models and pharmacological agents for selectively targeting nAChR have not been readily available until recently. For example, most *in vitro* model systems comprised of primary neurons are derived from the developing brain, and developing neurons fail to express and properly target functional nAChRs to synaptic junctions and extra-junctional locations (Srinivasan et al., 2011; Zambrano et al., 2015). Here, we leverage our observation that functional nAChR are expressed in acute hippocampal slice cultures derived from adult mice, and adopted our mouse model of acute intoxication with the OP diisopropylfluorophosphate (DFP) (Calsbeek et al., 2021) to transgenic mice with specific nAChR subunits genetically deleted to assess the role of nAChR in OP-induced SE.

## Results

### Primary neuronal/glial cell cultures from neonatal rodents express AChE but fail to target functional nAChR

Table 1 summarizes research conducted as part of the UC Davis CounterACT Center of Excellence whose goals included development of rapid throughput *in vitro* assays to identify novel therapeutic interventions against neuroactive threat agents. Several neuron/glia co-culture models derived from postnatal day (PND) 0-1 mouse or rat hippocampus or neocortex were investigated for their ability to form mature neuronal networks that expressed cell bound AChE and developed robust spontaneous calcium oscillations (SCO) patterns, which are regulated by the balance of excitation-inhibition mediated by the activity of ionotropic and metabotropic glutamatergic and GABAergic receptors (Cao et al., 2017; Cao et al., 2015). While SCO patterns in these primary neuronal models have been shown to be uniquely altered when exposed to pharmacological/toxicological chemicals that have distinct molecular mechanisms, they failed to respond to OPs (either DFP or paraoxon) at levels that inhibited ≥95% of their cellular AChE catalytic activity (Table 1). Cortical cultures established from embryonic day 18 mouse pups also failed to respond to OPs and compared to similar cultures derived from PND 0-1 mouse pups, expressed lower AChE activity at 14-17 days *in vitro*. Our attempts to culture primary neuron-glia co-cultures dissociated from the adult mouse cortex failed to produce sufficiently viable cultures that could be used for functional screening using the FLIPER Tetra to measure effects of chemical probes on SCOs. An important clue to explain this paradox came from the observation that primary neurons derived from the perinatal rodent brain developed robust responses to atropine but failed to develop responses to nicotine (Table 1), suggesting an obligatory role of one or more nAChR subtypes for eliciting responses to OPs.

**Table 1.**
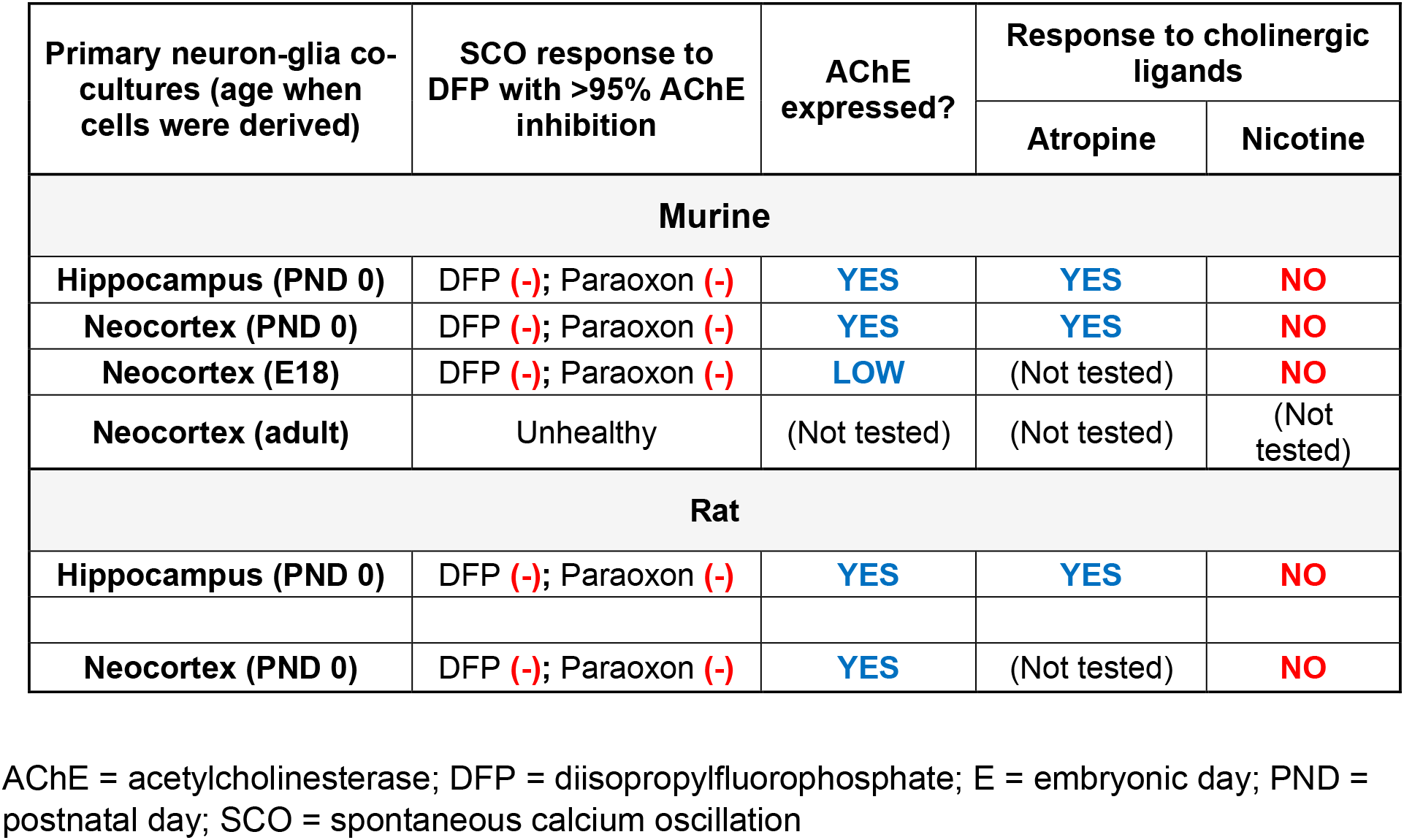
Common neuronal cell culture models express AChE but do not respond to OPs (DFP and paraoxon) or nicotine with altered spontaneous calcium oscillations (SCO)

| Primary neuron-glia co-cultures (age when cells were derived) | SCO response to DFP with >95% AChE inhibition | AChE expressed? | Response to cholinergic ligands |  |
| --- | --- | --- | --- | --- |
|  |  |  | Atropine | Nicotine |
| Murine |  |  |  |  |
| Hippocampus (PND 0) | DFP (-); Paraoxon (-) | YES | YES | NO |
| Neocortex (PND 0) | DFP (-); Paraoxon (-) | YES | YES | NO |
| Neocortex (E18) | DFP (-); Paraoxon (-) | LOW | (Not tested) | NO |
| Neocortex (adult) | Unhealthy | (Not tested) | (Not tested) | (Not tested) |
| Rat |  |  |  |  |
| Hippocampus (PND 0) | DFP (-); Paraoxon (-) | YES | YES | NO |
| Neocortex (PND 0) | DFP (-); Paraoxon (-) | YES | (Not tested) | NO |

AChE = acetylcholinesterase; DFP = diisopropylfluorophosphate; E = embryonic day; PND = postnatal day; SCO = spontaneous calcium oscillation

### DFP increased spontaneous electrical spike activity (ESA) proportionate to AChE inhibition in acute hippocampal slices prepared from adult mouse hippocampus

Mouse hippocampal slices were freshly prepared from adult male mice and randomly divided into two groups: one group exposed to either 3 or 20 µM DFP in artificial cerebrospinal fluid (aCSF) for 0-60 min and then frozen for later AChE determinations, and a second group that was mounted on perforated microelectrode arrays (pMEA) to record ESA before and after perfusion of 0, 3 or 20 µM DFP. Figure 1a portrayed the typical placement of a hippocampal slice on the pMEA. Once the slice was adhered to the pMEA and perfusion across the slice was adjusted, recording of electrical activity revealed spontaneous ESA in the absence of external electrical stimulation. Slices recorded on pMEA displayed spontaneous ESA that varied in basal frequency, although electrodes within or near CA1 typically produced the most active ESA (Figure 1b).

**Figure 1.**
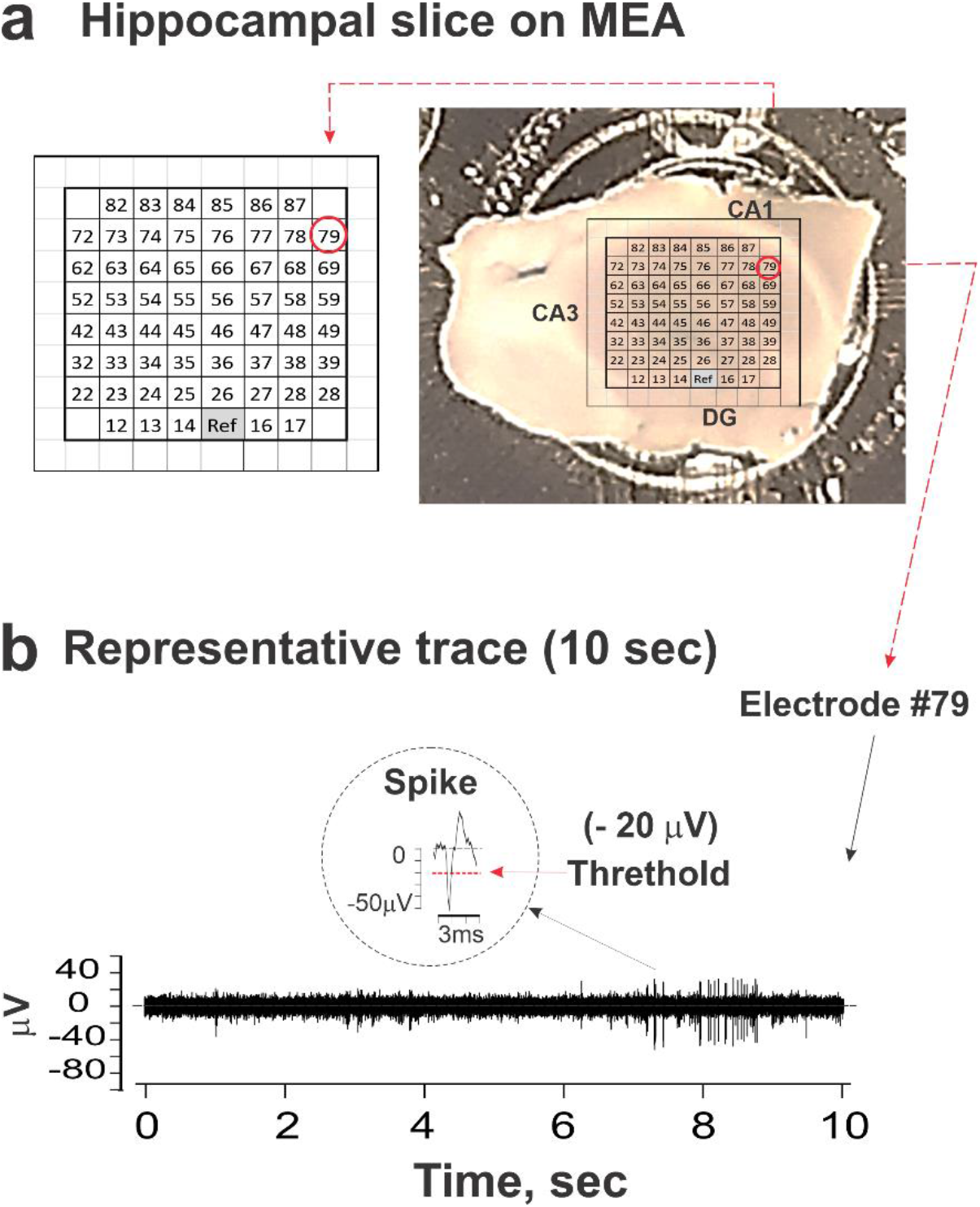
Recording of spontaneous network activity from mouse hippocampal slice on perforated microelectrode array (pMEA). Mouse hippocampal slices were firmly secured on pMEA using a dual-perfusion system. aCSF was perfused through the slice. **(a)** Although slices were consistently placed on the 60 electrode arrays to record from the entire hippocampus, including CA1, CA3 and DG, the slices were not placed stereotactically. **(b)** A representative 10- s recording of spontaneous electrical spike activity (ESA) from electrode #79. Each spike event was identified as a signal with a Falling Edge that exceeded the threshold of -20mV (red dashed line and arrow). The typical spike waveform of 3 ms is shown in the circle.

Figure 2a summarizes the relationship between AChE catalytic activity and the length of time the slices were exposed to DFP (3 and 20µM, filled-triangle and filled-circle, respectively). The rate with which DFP inhibited AChE was unexpectedly slow given the exposure paradigm consisted of through-slice perfusion of DFP. Addition of 3 and 20 µM DFP to the perfusate resulted in 90% inhibition of AChE activity relative to control aCSF within 37.4 ± 4.8 and 9.6 ± 2.3 min, respectively (Fig. 2b). ESA frequency increased with DFP exposure time and was tightly correlated with the degree of AChE inhibition, which was highly dependent on the DFP concentration (Fig 2a and b). Importantly, perfusion with DFP did not significantly increase ESA frequency until ≥80% of the slice AChE activity was inhibited and achieved peak ESA frequencies only when ≥95% AChE was inhibited (Fig. 2a).

**Figure 2.**
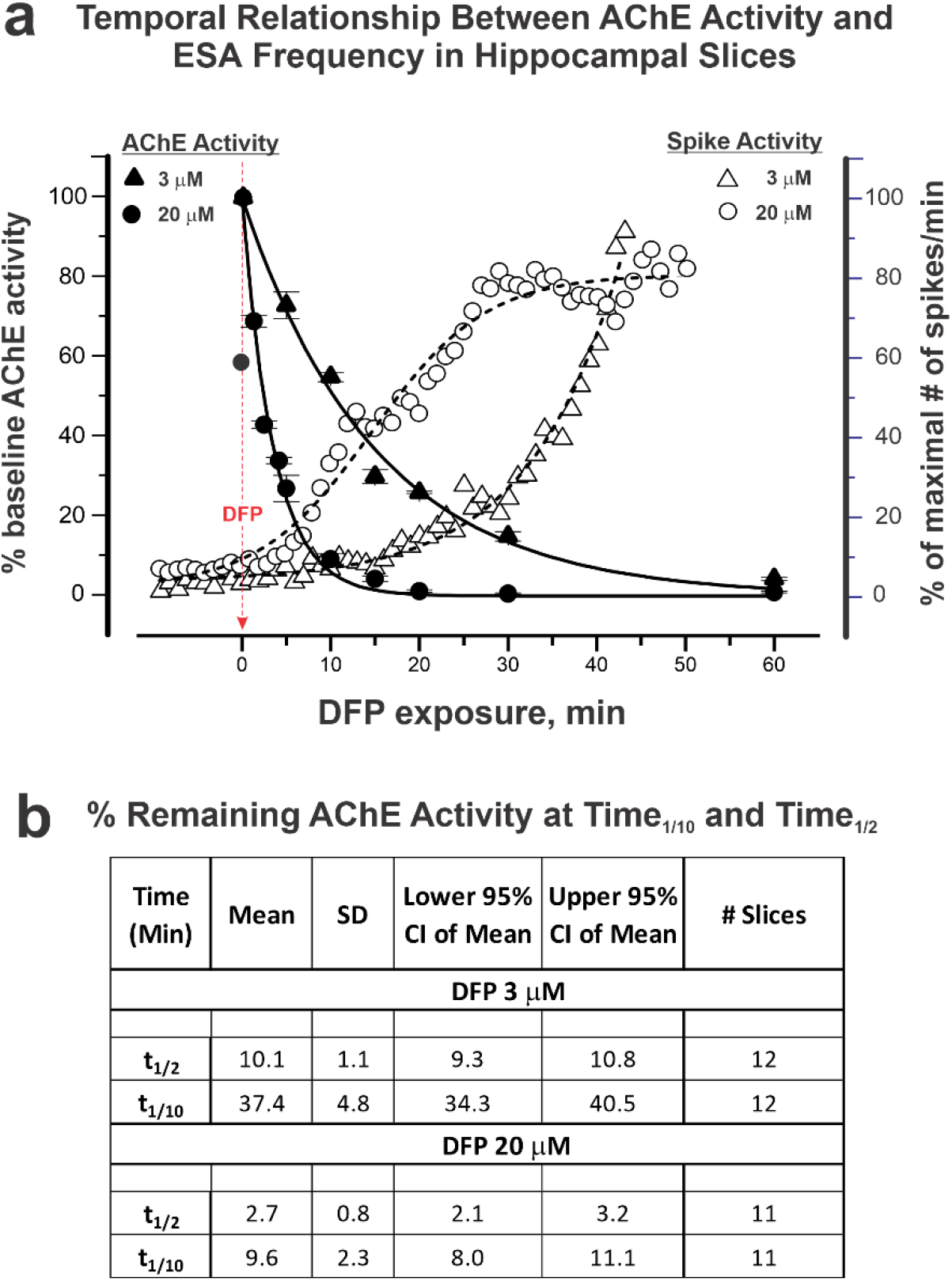
Temporal relationship between AChE activity and electrical spike activity (ESA) in hippocampal slices. AChE activity, measured using the Ellman assay, and ESA were assessed in acute hippocampal slices obtained from adult mice at varying times after addition of DFP at 3 or 20 µM. **(a)** AChE activity (left Y-axis; filled-triangles and circles) and ESA (right Y-axis; open-triangle and circles), both shown as a % of baseline activity prior to addition of DFP. The baseline AChE activity range was 0.41-0.50 mmole/min/mg. Each time point represents the mean value of n = 5 independent AChE preparations, each performed in triplicate (total 11-12 slices from N=28 mice). ESA frequency (right Y-axis) increased with exposure time and is presented as mean spikes/minute (% of maximum) from one representative slice exposed to 3 or 20 µM DFP (N=24 and N=17 active electrodes, respectively). **(b)** The time to achieve 50% and 90% inhibition of AChE (t_1/9_ and t_1/2_) were obtained by non-linear regression fitting using OriginLab.

Based on these results, all subsequent slice experiments were performed with 20 µM DFP in order to limit experiments to a total of 60 min (10 min baseline + 50 min with DFP perfusion). Figure 3a illustrates the experimental protocol used for further evaluating the temporal influences of DFP on ESA. The 60 min experiment was divided into six epochs, 10 min/epoch (Figure 3a). Epoch I recorded 10 min of baseline prior to addition of DFP to the aCSF perfusate and served as the basis for normalizing ESA for each active electrode. Epochs II-VI were recorded subsequent to addition of 20 µM DFP to the aCSF perfusate. Figure 3b shows a representative 10 s recording from each epoch and the overlay of all spike cutouts from these 10 sec recordings. Figure 3c shows the temporal changes in ESA frequency of all electrodes detecting activity in a single slice. Epochs II-VI from each active electrode were normalized to their respective baseline activity (epoch I) and reported as fold-change in spike rate over baseline (Fig. 3d). Statistical analyses of normalized changes in ESA frequency are summarized in Table 2. DFP (20 µM) produced a mean 9.1-fold increase in ESA frequency relative to baseline within the first 10 min (p=0.039, Table 2), whereas within the last epoch (epoch VI) DFP increased ESA frequency 67.1-93.6-fold (lower-upper 95% of mean, Table 2).

**Figure 3.**
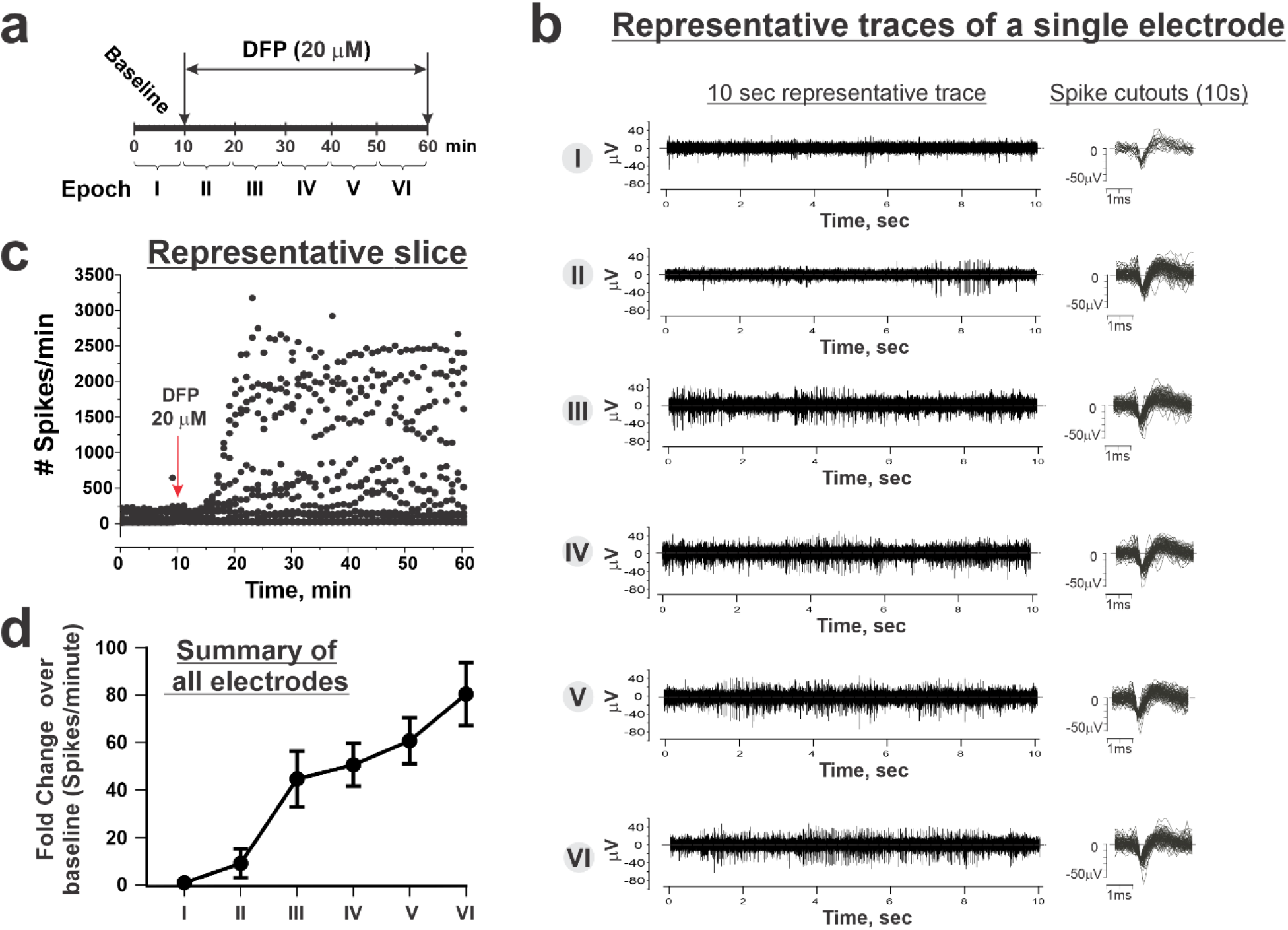
Temporal influence of DFP on hippocampal ESA. (a) Each slice recording was conducted over 60 min and analyses of ESA were subdivided into six consecutive epochs. The first 10 min served as the baseline and is defined as epoch I. DFP (20 µM) was added to aCSF perfusate after the first 10 min (indicated by the red arrow in panel c) with subsequent 10 min recordings defined as epoch II through VI. **(b)** Representative 10 sec recordings of ESA from a single electrode within each epoch. Spike cutouts of EPSP waveforms from each recording electrode were overlaid from each epoch. **(c)** ESA recorded from one slice from both active and inactive electrodes before and after addition of 20 µM DFP in the perfusate. **(d)** Mean ESA ± 95% confidence interval (CI) from all active electrodes from all slices (N=76 active electrodes from 10 slices obtained from 7 mice). The baseline recording (epoch I) from each active electrode was used to normalize that electrode’s DFP response. Statistical analyses of these data are presented in Table 2.

**Table 2.**
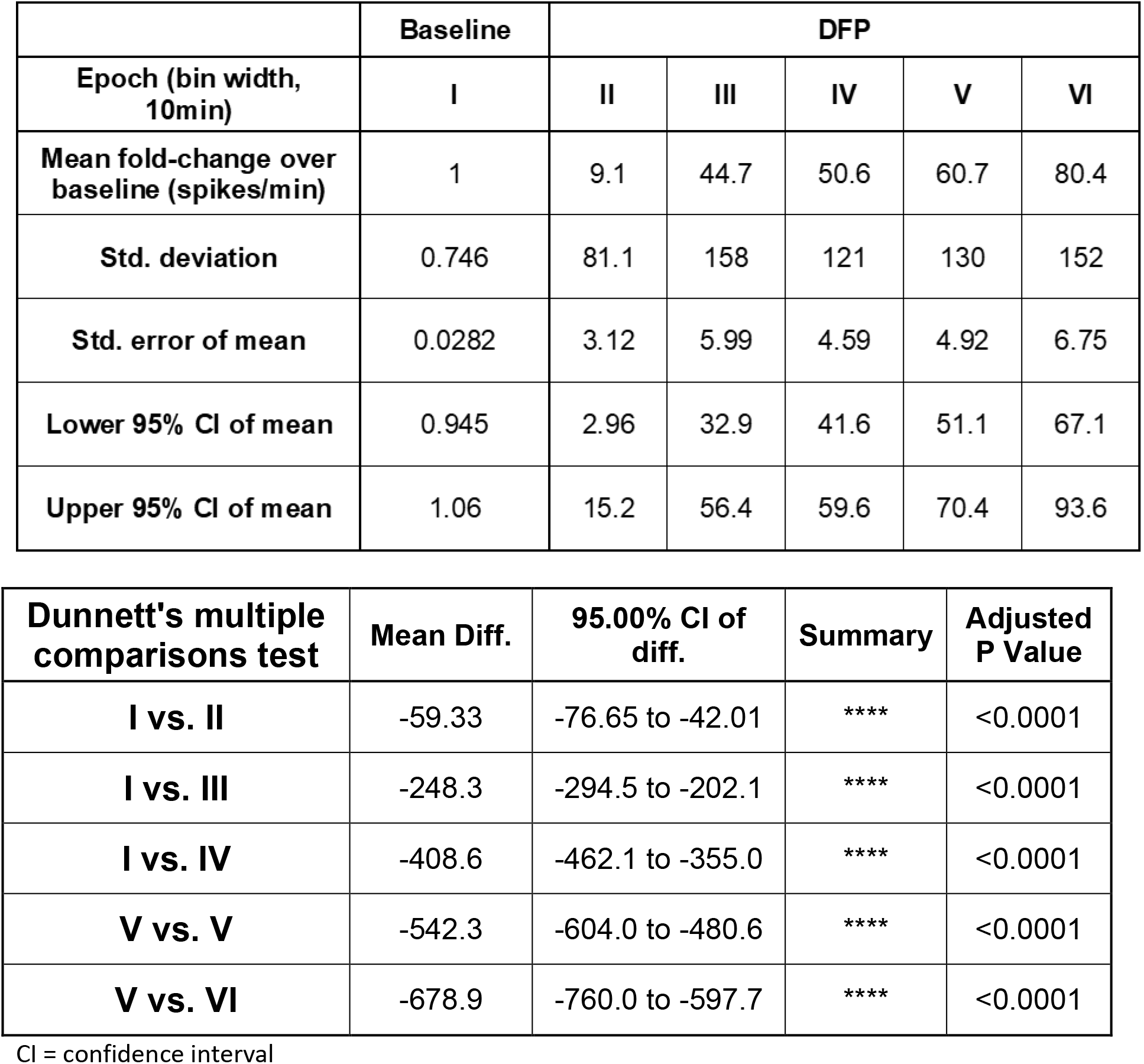
Statistical analysis of electrical spike activity (ESA; spikes/minute) represented as fold-change over baseline of same electrode.

|  | Baseline | DFP |  |  |  |  |
| --- | --- | --- | --- | --- | --- | --- |
| Epoch (bin width, 10min) | I | II | III | IV | V | VI |
| Mean fold-change over baseline (spikes/min) | 1 | 9.1 | 44.7 | 50.6 | 60.7 | 80.4 |
| Std. deviation | 0.746 | 81.1 | 158 | 121 | 130 | 152 |
| Std. error of mean | 0.0282 | 3.12 | 5.99 | 4.59 | 4.92 | 6.75 |
| Lower 95% CI of mean | 0.945 | 2.96 | 32.9 | 41.6 | 51.1 | 67.1 |
| Upper 95% CI of mean | 1.06 | 15.2 | 56.4 | 59.6 | 70.4 | 93.6 |

| Dunnett's multiple comparisons test | Mean Diff. | 95.00% CI of diff. | Summary | Adjusted P Value |
| --- | --- | --- | --- | --- |
| **I vs. II** | -59.33 | -76.65 to -42.01 | **** | <0.0001 |
| **I vs. III** | -248.3 | -294.5 to -202.1 | **** | <0.0001 |
| **I vs. IV** | -408.6 | -462.1 to -355.0 | **** | <0.0001 |
| **V vs. V** | -542.3 | -604.0 to -480.6 | **** | <0.0001 |
| **V vs. VI** | -678.9 | -760.0 to -597.7 | **** | <0.0001 |

CI = confidence interval

### Mecamylamine suppressed DFP-triggered ESA hyperexcitation

Mecamylamine (MEC), a non-selective noncompetitive antagonist of nAChR, was included in the aCSF perfusate in the absence or presence of DFP (Fig. 4a). Perfusion of MEC in the absence of DFP had no significant influence on ESA frequency compared to perfusion of aCSF alone (Figure 4b). Addition of MEC to the perfusate at the beginning of epoch II, 10 min prior to initiating DFP perfusion (20 µM) in epoch III, dampened ESA frequency compared to perfusion of DFP alone during epochs II-VI (Fig 4a and b). MEC suppressed DFP-triggered ESA frequencies 3.3-fold (p=0.064) during the first 10 min. In contrast, during subsequent epochs, MEC suppressed DFP-triggered ESA frequency to an even greater extent (Figure 4b and c).

**Figure 4.**
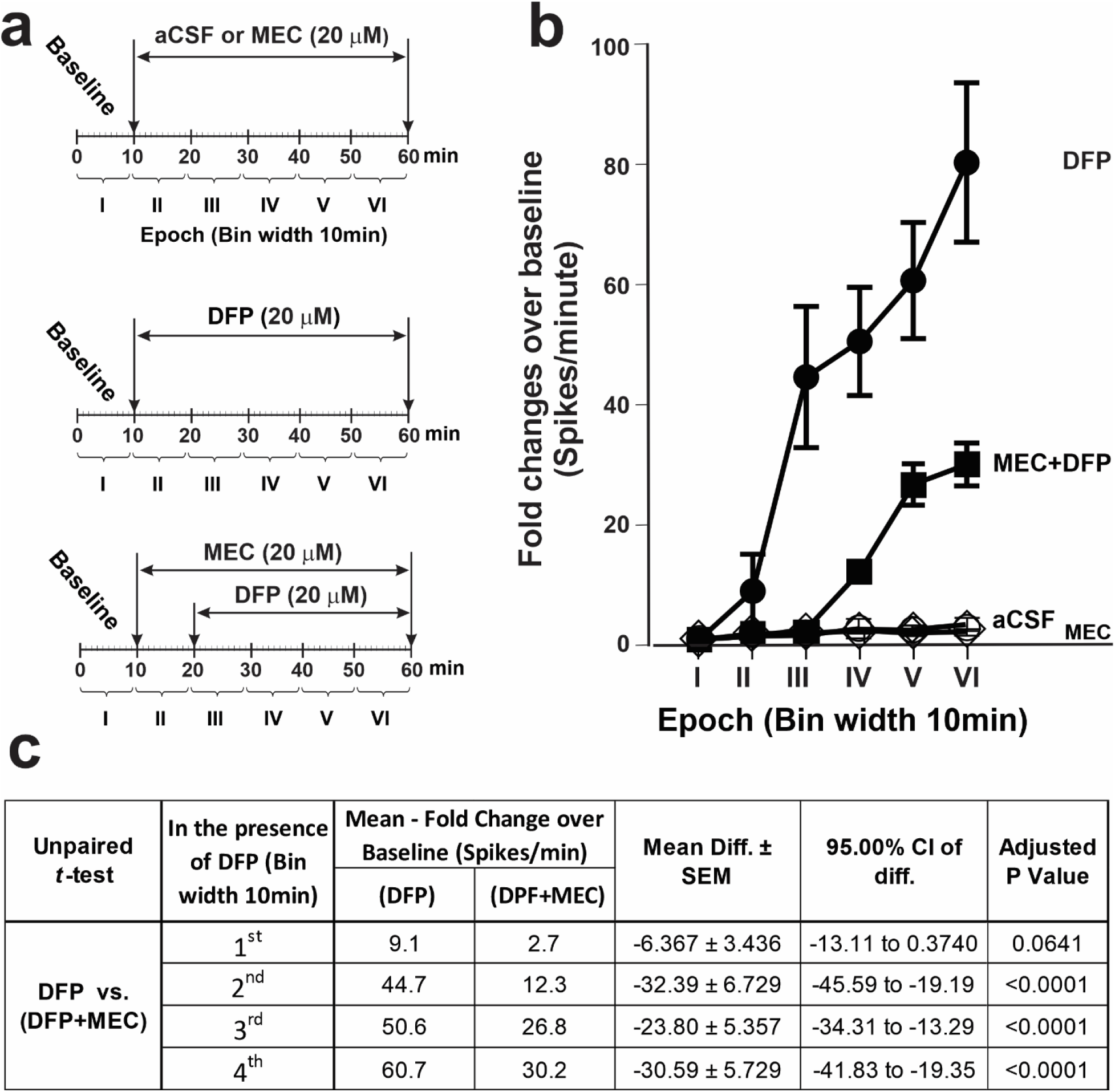
Mecamylamine (MEC) attenuated DFP-triggered hyperactivity of hippocampal slices. (a) Schematic illustration of various exposure paradigms assessed for effects on ESA. After a 10 min baseline recording, slices were perfused for an additional 50 min with either aCSF alone or aCSF to which MEC (20 µM) or DFP (20 µM) had been added. To determine whether MEC inhibited the effects of DFP on ESA, after 10 min of baseline recording, slices were perfused for 10 min with MEC (20 µM) and then with MEC and DFP (20 µM) for an additional 40 min. **(b)** Fold-changes in ESA relative to baseline within each of the six Epochs. Data shown as the mean ± 95% CI [aCSF (open circles), N=28 electrodes from 6 slices obtained 6 mice; the MEC alone group (open diamonds), N=23 electrodes from 4 slices obtained from 4 mice; DFP alone group (filled circles), N=76 electrodes from 10 slices obtained from 7 mice; MEC+DFP group (filled squares; N=56 electrodes from 6 slices obtained from 6 mice]. **(c)** Summary of the statistical analyses comparing ESA in the DFP group *vs*. the MEC+DFP group. Adjusted P value <0.05 was regarded as significant.

### Dihydro-β-erythroidine (DHβE) reversibly attenuated DFP-triggered increases in ESA

DHβE (20 µM) is a selective competitive antagonist of nAChR that multimerizes with the α4 nAChR subunit. When added to the perfusate 20 min after DFP, a time during which DFP-increased ESA frequency was obvious and accelerating (Figure 5a and b), DHβE significantly dampened, stopped or reversed DFP-mediated acceleration of ESA frequency measured from active electrodes compared with DFP alone (compare Fig. 4b to Fig 5b). DFP is an irreversible blocker of AChE, whereas DHβE is a reversible antagonist of α4 nAChR. Figure 5b shows that washout of both DFP and DHβE from the pMEA chamber for an additional 20 min quickly unmasked the irreversible nature of AChE inhibition by DFP, resulting in escalating ESA frequency. Comparing ESA frequencies recorded during perfusion with DFP and DHβE to those recorded after washout revealed a 2.5 ± 0.4-fold increase in ESA frequency, likely due to the enduring actions of DFP in the absence of DHβE (Fig 5c and d).

**Figure 5.**
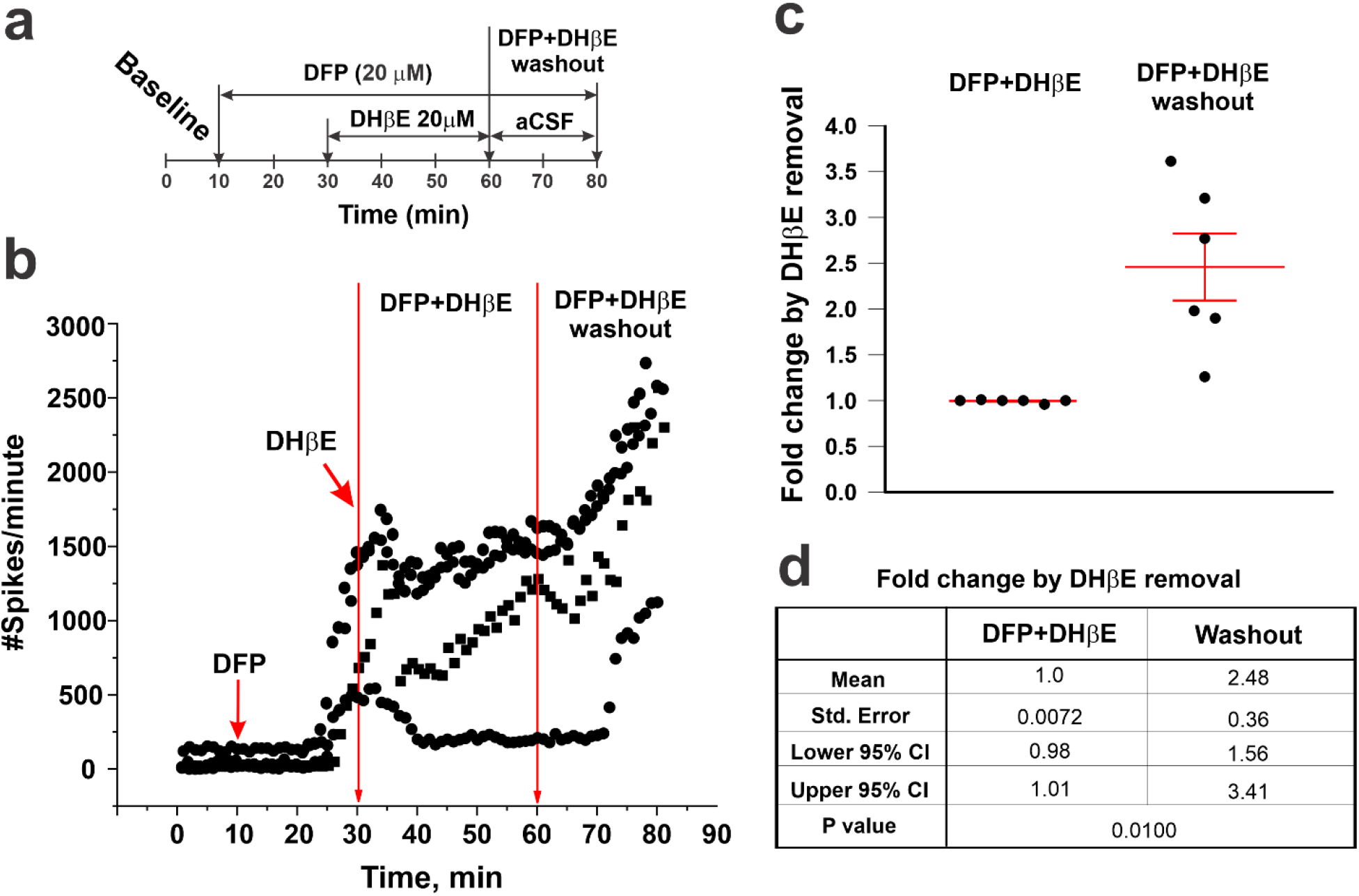
Selective α4 nAChR antagonism reversibly attenuated DFP-triggered hyperactivity. (a) Schematic of the exposure protocol: following a 10 min baseline recording, acute hippocampal slices were sequentially perfused with DFP (20 µM), the selective α4 nAChR antagonist, DHβE (20 µM), and then aCSF alone to remove DFP and DHβE. DHβE was introduced 20 min after the addition of DFP, at which time there was a significant increase in ESA. (b) ESA of three active electrodes from a representative slice. (c) Fold-change in mean ESA ± SEM after washout of DFP and DHβE (from 60-80 min) relative to ESA in the presence of both DFP and DHβE (from 30-60 min). Each dot represents a single mouse. (d) Summary statistics of data shown in panel c (N = 58 active electrodes from 6 independent slices from 6 mice).

### ESA responses to DFP are significantly attenuated in slices from α4 nAChR knockout (KO) mice

Among nAChR subtypes, pentameric assemblies with two α4 subunits were shown to be most abundantly expressed in the mammalian brain. Figure 6 shows that relative to slice from wildtype (WT) mice, DFP triggered significantly attenuated ESA responses in slices prepared from α4 nAChR KO mice. The epoch-wise comparisons of ESA frequency showed significantly lower ESA frequencies across all epochs perfused with DFP in α4 nAChR KO compared to WT slices (Figure 6c-d).

**Figure 6.**
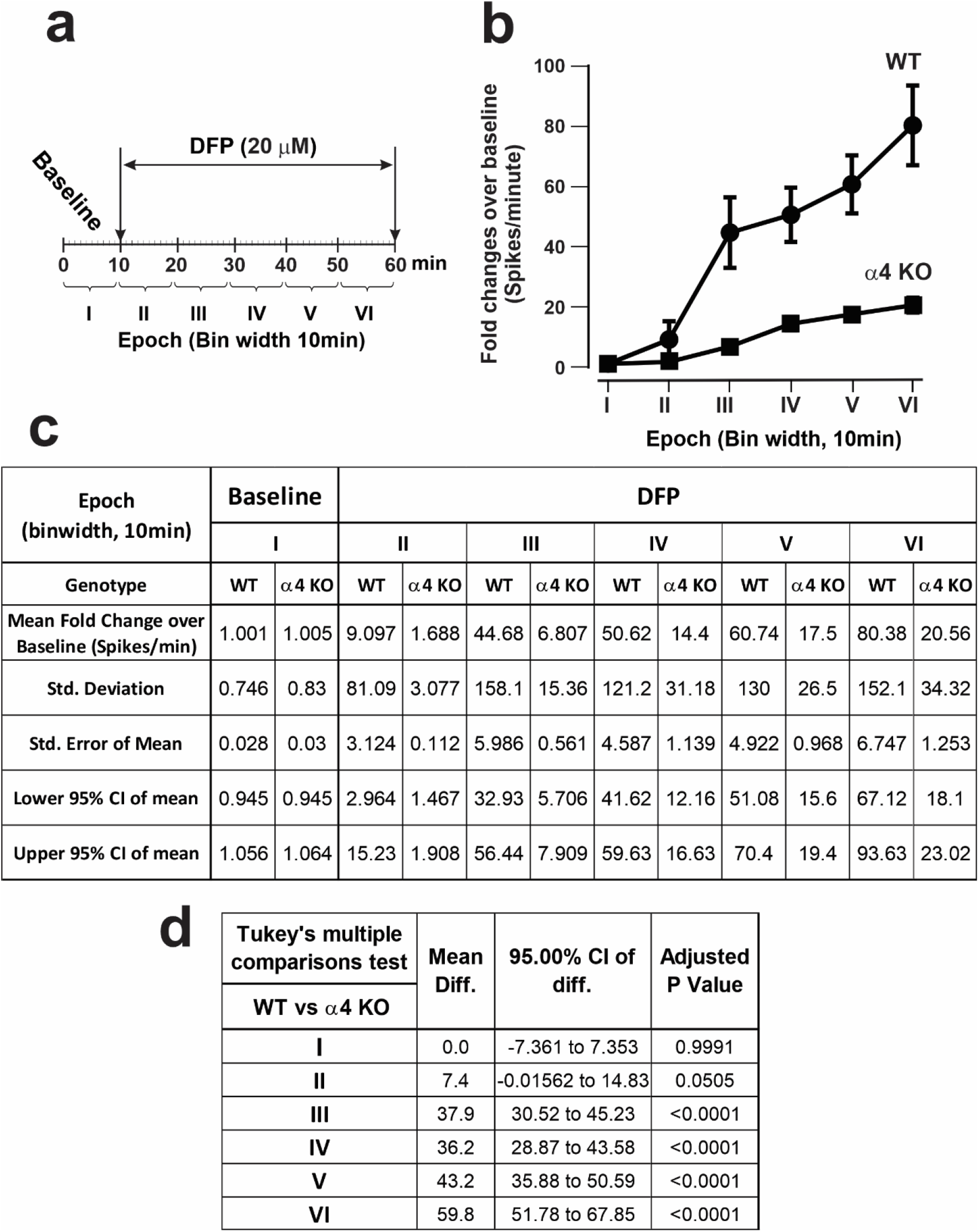
Greatly attenuated DFP responses in hippocampal slices from α4 nAChR KO mice. (a) Schematic of the experimental protocol in which DFP was added to hippocampal slices obtained from adult male wildtype (WT) or α4 nAChR KO mice. **(b)** The effect of genotype on DFP-triggered hyperactivity. Data are shown as the mean ESA ± 95% CI from epoch II through VI normalized to baseline (epoch I). **(c, d)** The descriptive statistics and Tukey’s multiple comparisons test. Adjusted P value <0.05 was regarded as significant. N=76 active electrodes WT mice N=7, N=10 slices with total. a4 KO mice N= 6; N=8 slices with total N=68 active electrodes.

### Pharmacologic antagonism or genetic KO of α4, but not α7, nAChR prevented DFP-induced EEG abnormalities

To confirm the physiological relevance of findings in acute hippocampal slices, we evaluated the effects of pharmacological antagonism or genetic deletion of nAChR on behavioral seizures and electroencephalographic (EEG) activity in a mouse model of acute DFP intoxication (Calsbeek et al., 2021). In our initial behavioral assessments, mice pretreated with either the nonspecific nAChR antagonist MEC or the α4-selective antagonist DHβE, and α4 KO mice displayed attenuated seizure activity compared to Veh/WT (Supplemental Fig. 1b), Moreover, broad nAChR receptor antagonism significantly attenuated seizure behavior when administered at 10 min, but not 40 min, after DFP administration (Supplemental Fig 1d). To confirm these behavioral data, subsequent experiments monitored seizure activity using electroencephalography (EEG). Representative EEG traces demonstrate that acute intoxication with DFP significantly altered EEG activity in a manner consistent with seizure activity (Fig. 7b and 8b). Quantitative analyses of these data confirmed that acute DFP intoxication triggered electrographic seizure activity. Specifically, in animals pretreated with vehicle, both the EEG amplitude as measured by the root mean squared (RMS) value, and the EEG spike rate were significantly increased within several minutes after DFP injection and both metrics remained consistently elevated relative to pre-DFP baseline values for hours post-DFP exposure (Fig. 7).

**Figure 7.**
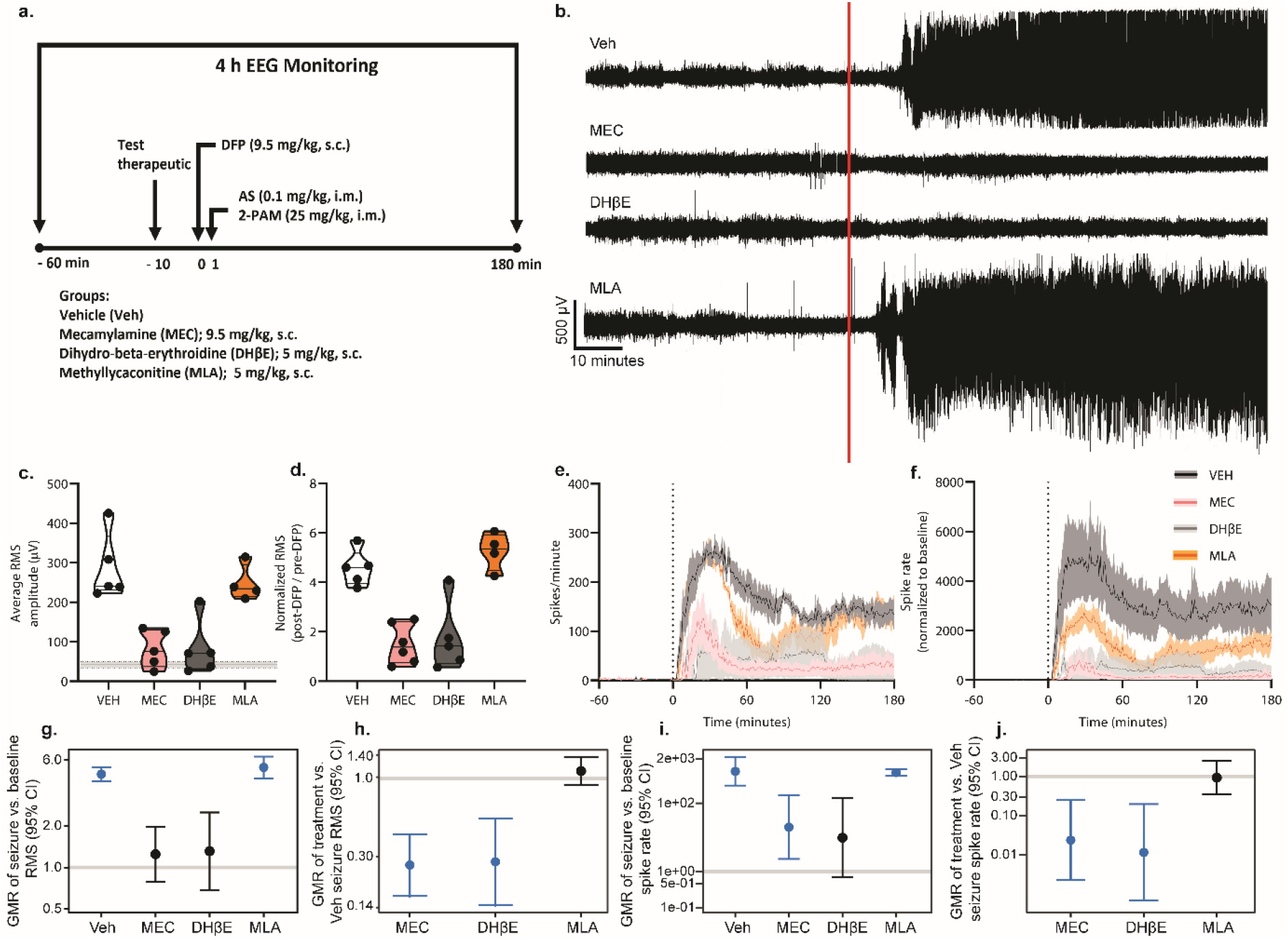
Pharmacologic antagonism of the α4 nAChR suppresses DFP-induced seizure activity. (a) Schematic illustrating study design. Adult male C57/BL6J mice were instrumented for EEG monitoring of seizure activity. On the day of experimentation, baseline EEG was recorded for 60 min prior to DFP intoxication. EEG monitoring continued for 180 min post-DFP injection. Mice were pretreated with vehicle (VEH), the nonselective nAChR antagonist mecamylamine (MEC), the α4-selective nAChR antagonist dihydro-β-erythroidine hydrobromide (DHβE), or the α7-selective nAChR antagonist methyllycaconitine citrate (MLA) 10 min prior to injection of DFP (9.5 mg/kg, s.c.), followed 1 min later by a combined injection of atropine sulfate (AS) (0.1 mg/kg, i.m.) and 2-pralidoxime (2-PAM) (25 mg/kg, i.m.). **(b)** Representative EEG traces for each experimental group. **(c)** Raw post-DFP RMS values in mice pretreated with either VEH or one of the nAChR antagonists. Pre-DFP baseline did not differ between groups and was therefore averaged across all groups and presented as a horizontal line ± SD. Data are presented as violin plots in which each point represents an individual animal, and the horizontal lines represent the minimum, quartile, median, and maximum values (n =4-6 per group). **(d)** Post-DFP RMS values normalized to pre-DFP baseline RMS in animals pretreated with vehicle or one of the pharmacologic antagonists of nAChR **(e)** Raw EEG spike rate and **(f)** spike rate normalized to baseline over the recording period in VEH (black) animals and animals treated with one of the pharmacologic antagonists of nAChR – MEC (pink), DHβE (gray), or MLA (orange). Data are presented as mean ± SEM (shaded areas) (n=4-6 per group). The vertical dotted line indicates the time of DFP administration. **(g)** Geometric mean ratio (GMR) (dot) of the mean normalized early-seizure RMS value for each group relative to pre-DFP baseline RMS with 95% confidence intervals (bars). The y-axis shown as log-scale. Confidence intervals that do not include 1 (the gray horizontal line) indicate a significant difference between pre- and post-DFP RMS values. (**h**) GMR of the mean normalized RMS values for each group relative to VEH normalized RMS. (**i**) GMR of the mean change in spike rate for each group between pre-DFP baseline and early-seizure periods. **(j)** GMR of the mean change in spike rate relative to VEH for each group between pre-DFP baseline and early-seizure periods.

**Figure 8.**
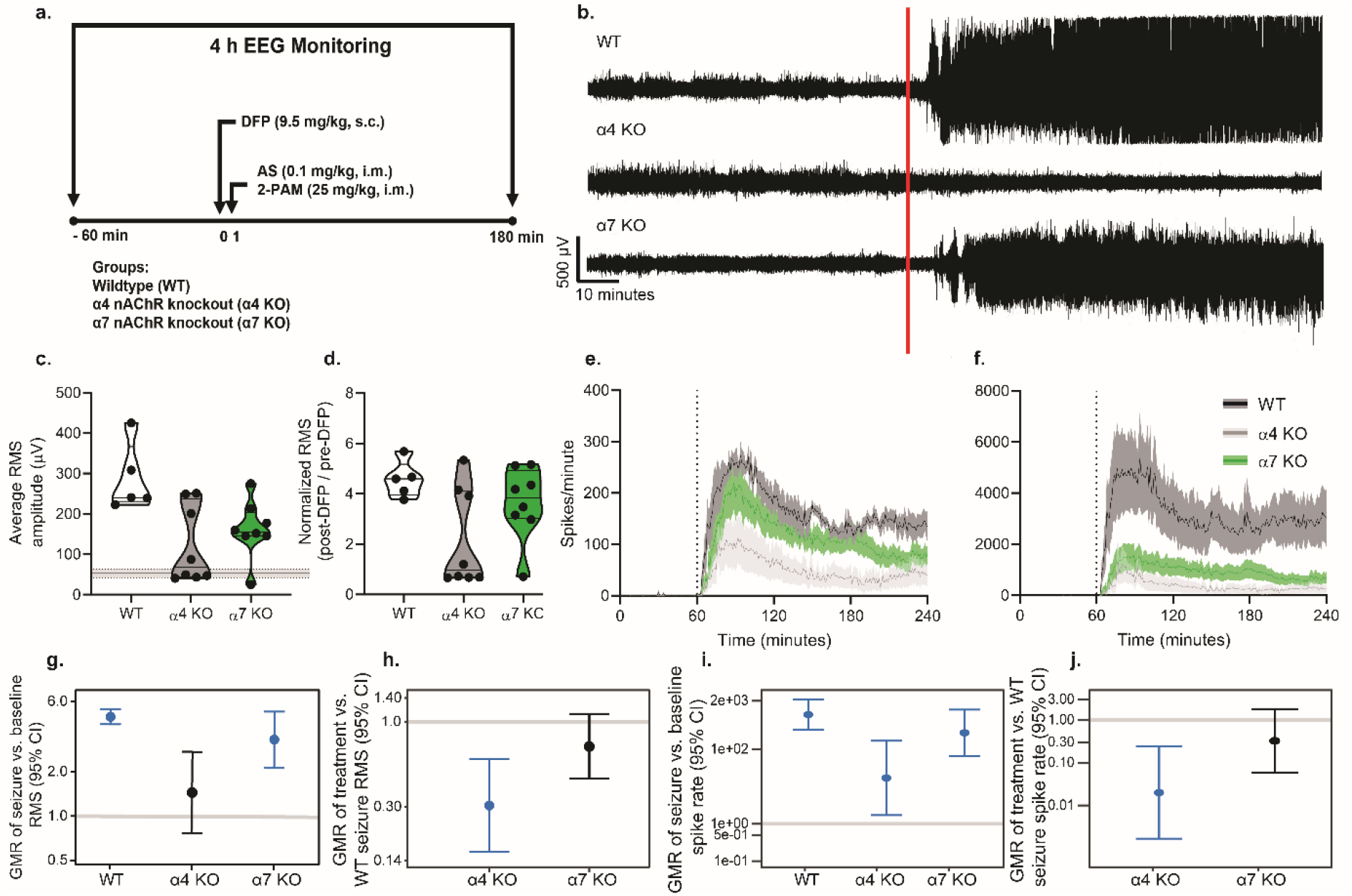
Genetic knockout of α4 nAChR attenuates DFP-induced seizures. (a) Schematic illustrating study design. Adult male wildtype (WT), α4 nAChR knockout (KO), and α7 nAChR KO C57/BL6J mice were instrumented for EEG monitoring of seizure activity. On the day of experimentation, baseline EEG was recorded for 60 min prior to DFP intoxication. EEG monitoring continued for 180 min post-DFP injection. WT animals are the same as VEH animals presented in Fig. 8 and were pretreated with vehicle (PBS) 10 min prior to injection of DFP. α4 and α7 KO animals did not receive any pretreatment. All animals were administered DFP (9.5 mg/kg, s.c.), followed 1 min later by a combined injection of atropine sulfate (AS) (0.1 mg/kg, i.m.) and 2-pralidoxime (2-PAM) (25 mg/kg, i.m.). **(b)** Representative EEG traces for each experimental group. **(c)** Raw post-DFP RMS values in WT, α4 KO, and α7 KO animals. Pre-DFP baseline did not differ between groups and was therefore averaged across all groups and presented as the horizontal line ± SD. Data are presented as violin plots in which each point represents an individual animal, and the horizontal lines represent the minimum, quartile, median, and maximum values (n =5-8 per group). **(d)** Post-DFP RMS values normalized to pre-DFP baseline RMS in WT, α4, and α7 KO animals. **(e)** Raw EEG spike rate and **(f)** spike rate normalized to baseline over the recording period in WT (black), α4 KO (gray), and α7 KO (green) animals. Data are presented as mean ± SEM (shaded areas) (n=5-8 per group). The vertical dotted line indicates the time of DFP administration. **(g)** Geometric mean ratio (GMR) (dot) of the mean normalized early-seizure RMS value for each group relative to pre-DFP baseline RMS with 95% confidence intervals (bars). The y-axis shown in log-scale. Confidence intervals that do not include 1 (the gray horizontal line) indicate a significant difference between pre- and post-DFP RMS values. **(h)** GMR of the mean normalized RMS values for each group relative to WT normalized RMS. **(i)** GMR of the mean change in spike rate for each group between pre-DFP baseline and early-seizure periods. **(j)** GMR of the mean change in spike rate relative to WT for each group between pre-DFP baseline and early-seizure periods.

Pharmacological antagonism (Fig. 7) or genetic deletion (Fig. 8) of α4, but not α7, nAChR prevented or significantly reduced DFP-induced seizure activity. WT mice pretreated with either the nonspecific nAChR antagonist MEC or the α4-selective antagonist DHβE (Fig. 7c), and α4 KO mice (Fig. 8c), exhibited post-DFP RMS values that were not significantly different from pre-DFP baseline values. In contrast, raw RMS values were significantly elevated post-DFP intoxication in WT mice pretreated with methyllycaconitine citrate (MLA, an α7-selective antagonist (Fig. 7c), as well as in α7 KO mice (Fig. 8c), and this relationship remained consistent after normalization of RMS to pre-DFP baseline values (Fig. 7d and Fig. 8d). Further analysis revealed that WT mice pretreated with vehicle (geometric mean ratio [GM]=4.7, 95% confidence interval [CI]=4.2-5.3, p<.001) or the α7-selective antagonist, MLA (GMR=5.3, 95% CI=4.4-6.3, p<.001) (Fig. 7g), as well as α7 KO mice (GMR=3.4, 95% CI=2.2-5.2, p<.001) (Fig. 8g), had RMS values during the early seizure period that were 3-5 times higher, on average, than their baseline values.

Raw and normalized EEG spike rate following DFP intoxication were also differentially altered by pharmacological (Fig. 7e,f), but not genetic (Fig. 8e,f), manipulation of nAChR signaling. All pharmacological treatment groups other than DHβE-pretreated WT mice exhibited a significant increase in spike rate during the first hour post-DFP exposure relative to baseline (Veh/WT: GMR=852.3, 95% CI: 330.1-2200.6, p<.001; MEC: GMR=20.1, 95% CI=2.4-171.4, p=.007; MLA: GMR=788.8, 95% CI=625.0-995.5, p<.001) (Fig 7i). Similarly, both genetic KO groups displayed significantly elevated spike rate during the first hour post-intoxication relative to baseline (α4 KO: GMR=17.0, 95% CI=1.7-169.2, p=.02; α7 KO: GMR=277.0, 95% CI=66.6- 1153.0, p<.001) (Fig. 8i).

We next compared the degree of electrographic response to acute DFP intoxication between experimental groups to determine which groups differed significantly from WT mice intoxicated with DFP in the absence of any pharmacological or genetic manipulation. WT mice pretreated with either MEC (GMR=0.3, 95% CI=0.2-0.4, p<.001) or DHβE (GMR=0.3, 95% CI=0.1-0.5, p<.001) (Fig. 7h) and α4 KO (GMR=0.3, 95% CI=0.2-0.6, p<.001), p=.003) mice (Fig. 8h) exhibited a 70% reduction in the post-DFP increase in RMS compared to Veh/WT controls. We did not observe significant differences between normalized RMS values in mice pretreated with MLA (Fig. 7h) or α7 KO mice (Fig. 8h) relative to Veh/WT controls.

The change in spike rate between baseline and post-DFP time points for each experimental group was also compared to Veh/WT controls intoxicated with DFP. The difference in spike rate between baseline and the early seizure period was significantly reduced in WT mice pretreated with MEC (GMR=0.02, 95% CI=0.002-0.25, p=.002) or DHβE (GMR=0.01, 95% CI=0.001-0.20, p=.002) (Fig. 7j) and in α4 KO (GMR=0.02, 95% CI=0.002-0.24, p=.002) mice (Fig. 8j) compared to Veh/WT controls. No significant differences in the rate of DFP-induced spiking were detected between controls and WT mice pretreated with MLA (Fig. 7j) or α7 KO animals (Fig. 8j).

Additionally, we evaluated weight loss following acute OP intoxication as a predictor of functional recovery (Rojas et al., 2021). Across all experimental groups there were no significant differences in body weight loss at either 4 or 24 h following DFP intoxication (Supplemental Fig. 2).

## Discussion

Previous studies have demonstrated that non-specific nicotinic antagonism suppresses parasympathetic symptoms and reduces mortality following acute OP intoxication (Harris and Stitcher, 1984; Hassel, 2006). While α4 (Fonck et al., 2003; Steinlein et al., 1995) and α7 (Orr-Urtreger et al., 1997; Stitzel et al., 1998) nAChRs modulate neuronal excitability, their involvement in OP-induced seizures has yet to be robustly investigated. Using complementary pharmacologic and genetic approaches in murine acute hippocampal slice preparations and an *in vivo* seizure mouse model, we found that α4, but not α7, nAChR signaling is involved in the induction of electrical spike activity and seizure activity following exposure to high levels of OPs.

With the mouse hippocampal slice preparation experiments, we found that inhibition of nAChR with the nonselective nAChR antagonist MEC or the α4 selective nAChR antagonist DHβE significantly attenuated DFP-triggered ESA when added either before (MEC) or within 10 min after DFP effects were elicited. Attenuated DFP-induced increases in ESA frequency were also observed in hippocampal slices prepared from α4 KO animals. These in vitro observations were predictive of *in vivo* effects in that electrographic metrics of DFP-induced seizure activity were significantly reduced by pretreatment with MEC or DHβE, and in α4 KO animals.

Our findings address the limited data regarding the role of nAChRs in models of acute OP intoxication. While early *in vivo* pharmacologic work emphasized the role of mAChR in OP-induced seizures, nAChR-sensitive soman-induced seizures have been reported earlier in the presence of a high, anti-convulsant dose of atropine sulfate (Shih et al., 1991). Such data hinted at a role for nAChR signaling in OP-induced seizures, but the lack of appropriate experimental tools complicated subsequent attempts to parse out the relative contribution of nAChRs *vs*. mAChRs in OP-induced seizure. The *in vivo* model used in these studies provided a low, peripherally-active dose of atropine sulfate to improve survival without robust central effects (Li et al., 2011). Using this model, we were able to demonstrate the role of nAChRs in OP-induced seizures independent of central mAChR antagonism.

The limited data available regarding the involvement of specific nAChR subtypes in models of acute OP intoxication has been due in part to the lack of cell culture models that appropriately target nAChRs to the neuronal cell surface, and thus, fail to respond to stimuli that activate nAChR signaling (Srinivasan et al., 2011; Zambrano et al., 2015). Here, we leveraged our observation that *in vitro* hippocampal slices prepared from adult mice were responsive to DFP, and used drugs that inhibited nAChR signaling with differing specificities to demonstrate that α4, but not α7, nAChRs significantly contribute to acute OP-induced hyperexcitability. Further, these observations were confirmed *in vivo*. While the role of the α4 nAChR subtype in OP-induced seizures has not been previously investigated, earlier studies identified the insensitivity of DFP-associated seizures to α7 nAChR signaling (Ferchmin et al., 2014), and the inefficacy of delayed broad nAChR antagonism or α7-selective antagonism in reducing soman-induced spiking in hippocampal slice preparations (Harrison et al., 2004). Our findings, paired with these observations, confirm the α7-independence of OP-induced spiking. Additionally, the ineffectiveness of delayed nAChR antagonism is consistent with the timeline of the cholinergic-glutamatergic transition required for maintenance of OP-induced seizures (McDonough and Shih, 1997).

In non-OP models, it is established that nAChR activation can induce seizures; however, the role of receptor subtypes in seizure generation is not well understood. Nicotine-induced seizures are blocked by broad nAChR antagonism (Damaj et al., 1999), but there are discrepant reports regarding the involvement of α4 *vs*. α7 nAChRs. Peripheral and central administration of α7 selective antagonists, including MLA or α-nudicauline, were shown to attenuate nicotine-induced seizures, but because their mitigating effects were significantly less potent than those produced by MEC and occurred in a dose range known to affect nicotinic responses not mediated by α7, the authors concluded that involvement of α4β2-receptor subtypes could not be ruled out (Damaj et al., 1999). The authors also concluded that the α7 nAChR subtype does not play a major role in initiating nicotine-induced seizures, a conclusion that is consistent with our observations of the lack of effect on DFP-induced seizures with pharmacological antagonism or genetic deletion of the α7 nAChR. Homozygous α5 nAChR subunit KO mice are highly resistant to nicotine-triggered seizures in the presence or absence of the β4 nAChR subunit (Kedmi et al., 2004; Salas et al., 2003), whereas a genetic model of α4 hypersensitivity exhibited increased susceptibility to nicotine-induced seizures (Fonck et al., 2003). Previous studies of the α4 nAChR subunit KO mice indicated that compared to WT they did not exhibit spontaneous seizures but did exhibit heightened anxiety and an altered time course in developing behavioral topographies in response to non-seizurogenic doses of nicotine (Ross et al. 2000). Furthermore, α4 nAChR subunit KO mice had a greater sensitivity to the GABA receptor blockers, pentylenetetrazol and bicuculline, and to a lesser extent, the glutamate receptor agonist kainite (Wong et al., 2002). In contrast, α4 KO mice were significantly protected from 4-aminopyridine-induced major motor seizures and death compared to WT (Wong et al., 2002). Collectively, these results point to the importance of α4 and α5 nAChR subtypes in modulating sensitivity to seizure induction by nicotine. However, whether either subtype is essential for OP-triggered seizure induction, which is characterized by elevated synaptic acetylcholine (ACh) (Lallement et al., 1992), has not been directly tested. While ACh is known to mediate seizure generation in epileptic models (Zimmerman et al., 2008) (Maslarova et al., 2013), this relationship has not been comprehensively evaluated in naïve animals. Similarly, in slice preparations, exogenous ACh exerted excitatory effects (Dodd et al., 1981; Vidal and Changeux, 1989; Wheal and Miller, 1980), but the capacity for excessive synaptic ACh to induce seizures and the role of particular AChR subtypes in ictal activity, has not been rigorously investigated. Our findings address this data gap, providing the first definitive evidence that α4 nAChR subtypes are necessary for neuronal hyperexcitability induced by cholinergic receptor hyperstimulation, and are involved in OP-induced seizures *in vivo*.

We observed that α4 antagonism post-DFP incubation attenuated and occasionally reversed DFP-triggered ESA hyperactivity in individual hippocampal slices, possibly the result of rapid engagement of additional receptor subtypes or neurotransmitter systems post-OP exposure. Our *in vivo* observation that MEC administered 10 min, but not 40 min, post-DFP exposure attenuated seizure activity (Supplemental Fig. 1d), further suggested rapid activation of non-α4 nAChR subunits following acute OP intoxication. The ability of MEC to inhibit seizure activity when administered shortly after DFP exposure may reflect the fact that glutamatergic systems are engaged within 5 min of acute OP intoxication (McDonough and Shih, 1997), and high concentrations of MEC (∼100 µM) inhibit NMDA-mediated currents (O’Dell and Christensen, 1988). It has previously been demonstrated that MEC administered at 5 mg/kg, i.v. resulted in brain concentration of MEC of ∼ 42 µM (Debruyne et al., 2003), thus, it is possible that our dosing paradigm achieved brain concentrations of MEC high enough to inhibit both nAChR and NMDA receptors. Additional investigation is needed to fully resolve the temporal relationship between acute OP intoxication and activation of distinct neurotransmitter systems/receptor subtypes in the brain.

As expected with intrinsically different models (acute hippocampal slice preparations *vs*. *in vivo* cortical EEG monitoring), there was some quantitative disparity in the effect of pharmacologic and genetic modulation of nAChR signaling on the response to DFP. For example, we observed more pronounced suppression of DFP-induced seizure activity/hyperexcitability *in vivo* compared to slice preparations. Previous publications have described the necessity of an antimuscarinic for efficacy of nAChR antagonism against non-electrographic aspects of OP-associated toxicity (Chiou et al., 1986; Fleisher et al., 1970), suggesting that our *in vivo* use of atropine sulfate and 2-pralidoxime may influence response to DFP exposure. Additionally, OP-induced seizures are thought to originate external to the hippocampus (McDonough and Shih, 1997), and while acute hippocampal slice preparations replicate many features of the intact nervous system, they lack afferent input (Lein et al., 2011). Such differences in circuitry composition and/or degree of circuit engagement in hippocampal slice spike activity versus *in vivo* monitoring may underlie differences in our models.

*In vivo* seizure monitoring revealed a bimodal effect of α4 nAChR KO on seizure induction. While the majority of α4 nAChR KO animals did not display electrophysiologic shifts characteristic of seizure following acute DFP intoxication, the initial response of a subpopulation of the KO animals to DFP was similar to that of WT animals. The same bimodal distribution was not seen with pharmacologic inhibition of α4 nAChR. There is limited evidence that global α4 KO produces compensatory shifts in cholinergic signaling (Klink et al., 2001), which could variably influence response to acute OP intoxication. Pharmacospecificity may underlie differences in response to DHβE treatment and α4 KO; DHβE has been shown to inhibit α3 nAChRs under certain conditions (Harvey et al., 1996). While our data suggest α4 signaling as a primary driver of OP-induced seizures, α3 nAChRs are known to contribute to nicotine-associated seizures (Salas et al., 2004). It is possible that α3 nAChRs may be involved in OP spiking, although to a limited degree. The involvement of additional receptor subtypes in OP-induced spiking is further suggested by our observation of significantly attenuated, but still elevated, electrical spike rate immediately following DFP intoxication in α4 KO animals compared to WT animals. Thus, while our pharmacologic data demonstrated clear evidence that α4 nAChR signaling is involved in DFP-induced seizures, α4 KO does not prevent all OP-associated hyperexcitability. It is likely that incomplete suppression of DFP-induced spiking in α4 KO animals underlies the limited, but still evident, behavioral seizures in these animals (Supplemental Fig. 1c). Additional investigation will be needed to clearly define the role of α3 and additional nAChRs in OP-mediated seizures.

Our findings are important to advance the development of anti-seizure therapeutics, particularly prophylactics for individuals at high risk of acute OP intoxication. Historically, such individuals have received pyridostigmine bromide (PB), a reversible AChE inhibitor, to mitigate OP-associated toxicity. However, PB use has been tied to a number of adverse health outcomes (Chao et al., 2014; McCauley et al., 2001), highlighting the need for prophylactic alternatives. While we observed comparable anti-seizure efficacy with broad nAChR and α4-selective antagonism, non-selective nAChR inhibitor ganglionic blockers can produce concerning autonomic consequences (Nies, 1975). In contrast, our data identified α4 nAChRs as a feasible target for future prophylactic development warranting scientific investment.

In summary, we offer compelling evidence for the necessary role of α4 nAChR activation in seizure induction following acute OP intoxication. We demonstrated that OP-induced seizures are sensitive to genetic or pharmacologic inhibition of α4, but not α7, nAChR and that these observations are consistent between *in vitro* and *in vivo* assessments. These findings advance our understanding of the nicotinic cholinergic system in seizure activity and have implications for therapeutic development.

## Materials and Methods

### Materials

DFP and all test pharmaceuticals were obtained from the same sources for *in vitro* and *in vivo* studies. DFP was purchased from Sigma (St. Louis, MO, USA) and stored at -80°C. DFP purity ≥ 90% was confirmed by nuclear magnetic resonance as previously described (Gao et al., 2016). Atropine sulfate (AS), 2-pralidoxime (2-PAM) and paraoxon were purchased from Sigma and stored at room temperature of -20 °C. Manufacturer certificates of analysis indicated the purity of AS (lot #BCBM6966V) to be > 97%, 2-PAM (lot #MKCG3184) to be > 99%, and paraoxon (#MFCD00007316) to be 99.9%. Nicotine was purchased from Fisher Scientific (Fisher Scientific; Fair Lawn, NJ 07410). Manufacturer certificates of analysis indicated the purity of nicotine (#AC181420050) was >99%. Mecamylamine hydrochloride was purchased from AK Scientific (Union City, CA, USA). Dihydro-β-erythroidine hydrobromide (DHβE) and methyllycaconitine citrate (MLA) were purchased from Tocris (Bristol, UK). Manufacturer certificates of analysis indicated the purity of MEC (lot #TC23203) to be > 98%; DHβE (batch #11A/249912), > 98%; and MLA (batch #23A/251394), >98%.

### Animals

All experiments involving animals were performed in accordance with the National Institutes of Health Guide for the Care and Use of Laboratory Animals following protocols approved by the University of California, Davis (UC Davis) Institutional Animal Care and Use Committee.

Adult male WT C57BL/6J mice (8-10 weeks old) were obtained from The Jackson Laboratory (Sacramento, CA, USA) and allowed to acclimate for at least 7 d prior to surgical instrumentation for EEG measurement or experimentation. Three breeding pairs of heterozygous α4-nAChR global knockout (KO) mice on a C57BL/6J background (Ross et al., 2000) provided by Henry A Lester, PhD of the California Institute of Technology (Pasadena, CA, USA) and three breeding pairs of homozygous α7-nAChR global KO mice on a C57BL/6J background (Orr-Urtreger et al., 1997) (RRID #003232) obtained from The Jackson Laboratory (Sacramento, CA, USA) were bred at UC Davis to generate homozygous α4-nAChR KO mice and homozygous α7-nAChR KO mice that were used as adults for experimentation. The genotype of each animal used for experimentation was confirmed upon weaning at PND 21 using the primers described in Table 3. Additional details for genotyping the α4-nAChR KO animals are provided in the supplemental material. Homozygous α7-nAChR KO animals were genotyped using the protocol provided by the Jackson Laboratory (https://www.jax.org/Protocol?stockNumber=003232&protocolID=28408).

**Table 3.**
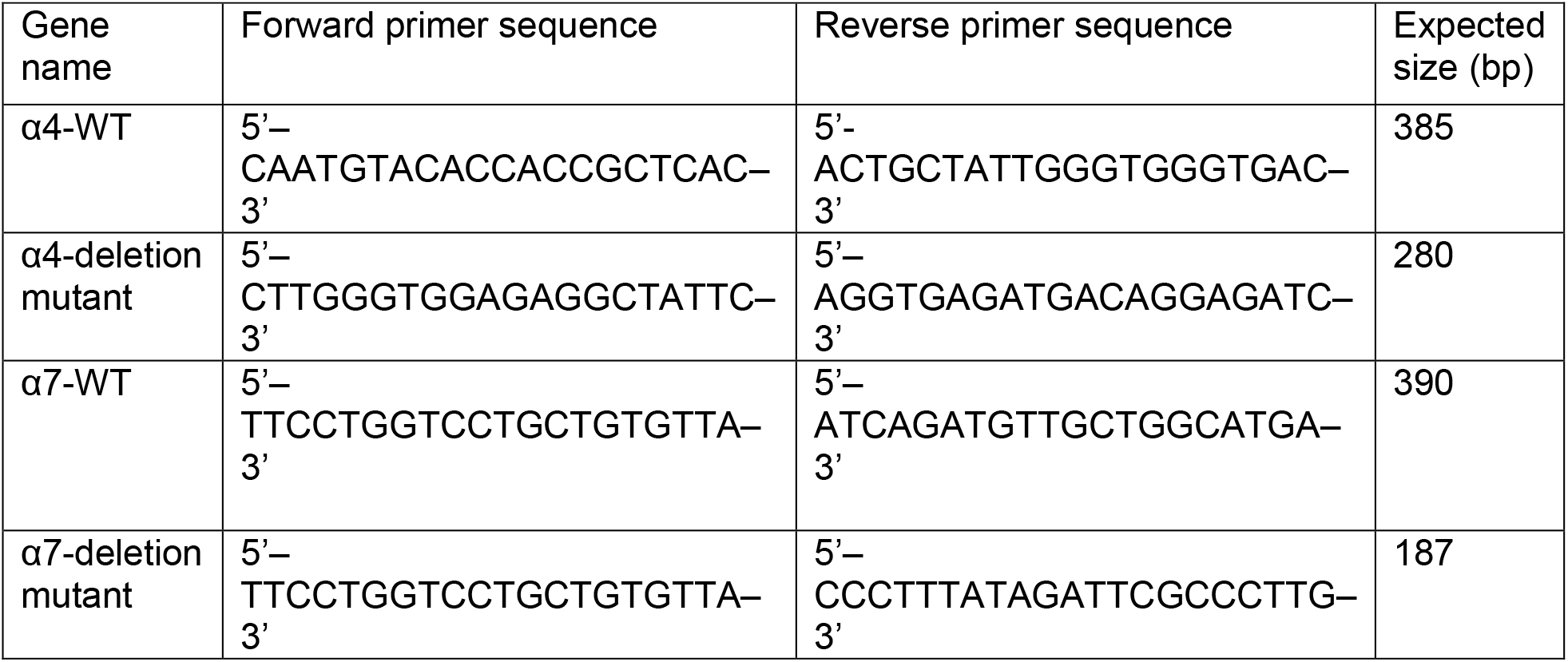
Genotyping sequences used in the study.

All weaned and adult mice were group housed (up to 4 mice per cage) in standard plastic cages in facilities fully accredited by AAALAC International under controlled environmental conditions (22 ± 2°C, 40-50% humidity, 12:12 light-dark cycle). Animals were provided standard rodent chow (LabDiet #5058) and tap water *ad libitum*.

### Primary neuron-glia co-cultures

Briefly, hippocampal or cortical neuronal/glial cultures were plated from C57BL/6J mice at embryonic day 18 (E18) or PND 30-45 and from Sprague Dawley rat pups at PND 0–1. For E18 and PND 0-1 cultures, brains from male and female pups were combined, whereas for adult cultures, only male mice were used using an established protocol (Kretz et al 2007). Dissociated brain tissues were plated onto poly-L-lysine-coated, clear-bottomed, black walled 96-well imaging plates (BD Bioscience, Franklin Lakes, New Jersey) at a density of 75,000 cells/well in complete Neurobasal medium (ThermoFisher, Waltham, Massachusetts) supplemented with 0.5 mM L-glutamine, GS21 supplement (Cat No. GSM-3100; MTI-GlobalStem Products, ThermoFisher, Gaithersburg, Maryland), 10 mM HEPES, and 5% fetal bovine serum (FBS) (Atlanta Biologicals, Norcross, Georgia). FBS concentration was reduced to 2.5% in the first day and halved at each change of media. For microelectrode array assays, 110,000 cells/well were seeded in a 12-well Maestro plate (Axion BioSystems, Atlanta, Georgia). Cytosine arabinoside (5 μM; Sigma Aldrich, St Louis, Missouri) was added to all cultures at 2 days *in vitro* and half of the media was changed every 3–4 days (the cytosine arabinoside concentration halved at each change of media). Neurons were incubated at 37°C with 5% CO_2_ and 95% humidity for 7–16 days.

### Measurement of Ca^2+^ in primary neuron-glia co-cultures

In primary neuron-glia co-cultures, SCOs were measured using the FLIPR TETRA plate reader system as previously described (Antrobus et al., 2021; Cao et al., 2017; Cao et al., 2015; Feng et al., 2017; Kim et al., 2023; Zheng et al., 2019). The FLIPR TETRA plate reader platform continuously measures and records intracellular Ca^2+^ transients from 96-wells, simultaneously, and responses from each well were compared to their own baseline activity. The culture medium was replaced with prewarmed (37°C) Locke’s buffer composed of (in mM): NaCl 154, KCl 5.6, CaCl_2_ 2.3, MgCl_2_ 1, HEPES 8.6, glucose 5.6, glycine 0.0001, pH 7.4, 0.0005 Ca^2+^ fluorescence dye Fluo-4-AM (Sigma-Aldrich) and 0.5 mg/ml bovine serum albumin fraction V (Fisher Scientific). Plates were returned to the culture incubator for 1 h at 37°C. Next, cells were gently washed with Locke’s buffer 3 times and permitted to equilibrate for 10–15 min. Baseline SCO recordings were taken for 10 min. Then the programmable 96-channel pipetting robotic system added either control vehicle or test chemicals, either individually or combined (e.g.., ACh+DFP), to the wells. Changes in SCO patterns (frequency and amplitude) were monitored for an additional 20 min. ScreenWorks Peak Pro Software (Version 3.2; San Jose, California) was used to normalize the magnitude of SCOs based on (*F*_max_ – *F*_min_)/*F*_min_, where *F*_max_ was the peak for each spike, and *F*_min_ was the baseline just prior to the spike. Kinetic Reduction Configuration was used to score the amplitude of SCOs. Total SCO frequency was scored using Peak Frequency (BPM) with Configure Peak Detection with the following settings: smooth width 1, fit width 3, slope threshold 0.01, amplitude threshold dynamic 7. Total SCO amplitude was scored using Average Peak Amplitude with Configure Peak Detection with the following settings: smooth width 1, fit width 3, slope threshold 0.01, and amplitude threshold dynamic 7.

### Preparation of acute hippocampal slices

Whole mouse brains dissected from adult male mice on PND 60-80 immediately after cervical dislocation were placed in a petri dish with ice-cold sucrose cutting solution consisting of (in mM): sucrose 150, NaCl 40, KCl 4, NaH_2_PO_4_H_2_O 1.25, CaCl_2_2H_2_O 0.5, MgCl_2_6H_2_O 7, glucose 10 and NaHCO_3_ 26. Each brain was affixed with glue to the plate of a vibrating blade microtome (VT1000S; Leica, Bannockburn, IL). The plate was submerged and maintained in the ice-cold, carbogen-gassed (95% O_2_+5%CO_2_, medical grade) sucrose cutting solution. Horizontal cutting thickness was set at 400 µm. Two to three slices from the medial hippocampus of each hemisphere were transferred to a recovery chamber (UNICO Incubator, L-CU60; Amazon.com; USA) containing aCSF (composed of, in mM: NaCl 125, KCl 3.5, NaH_2_PO_4_H_2_O 1.2, CaCl_2_2H_2_O 2.4, MgCl_2_6H_2_O 1.3, glucose 25 and NaHCO_3_ 26) at 32°C constantly bubbled with carbogen. After a 60 min stabilization period, slices were transferred onto pMEA (60pMEA200/30iR-Ti; Multichannel Systems GmbH, Reutlingen, Germany).

### Measurement of AChE activity in mouse neuron/glial cell cultures and hippocampal slices

AChE activity was measured in hippocampal slices setup using the same protocol described above for setting up slices on pMEA. After recovery, hippocampal slices were grouped (4-6 slices/group/mouse) and transferred to a container with DFP at 0 (aCSF only as control), 3 or 20 µM in aCSF/carbogen for 0, 5, 10, 15, 20, 30, 45 or 60 min at 32 °C. At the end of each incubation time, slices were transferred to cryotubes and stored in -80°C freezer until AChE determinations were performed.

Hippocampal slices or neuronal/glial cell cultures were homogenized in phosphate buffered saline (PBS; NaCl 137 mM, KCl 2.7 mM, Na2HPO4 10 mM, KH2PO4 1.8 mM, pH 7.4) supplemented with 1 mM EDTA (pH 8.0) using glass-Teflon homogenizer. The homogenate was centrifuged at 700 x g at 4°C for 10 min. The supernatant was collected and centrifuged at 100,000 x g for 60 min. The pellet was resuspended in PBS/EDTA buffer and the protein concentration of the homogenate measured using the Pierce BCA protein assay kit (Thermo Scientific, USA). AChE activity was measured using the Ellman assay using acetylthiocholine as substrate and the indicator 5,5′-dithiobis-2-nitrobenzoic acid (DTNB) mixed with 10 µg hippocampal lysate protein per well in 96-well plates (Falcon) (Ellman et al., 1961). The O.D./min was continuously recorded at 405 nm over 30 min to measure the hydrolytic product 5-mercapto-2-nitrobenzoic acid using a Synergy H1 microplate reader (Biotek; VT, USA). O.D./min was converted to catalytic rate using the Beer-Lambert law:

AChE activity (μmole/min) = {Slope (OD/min) × 2.5 x 10^-4^ (liter) × 10^6^ (μmole/mole)} {14,150 [liter/(mole × cm)] × 0.125 cm}.

GraphPad Prism software (Version 8; San Diego, California) was used for data analysis and graphing. The values of IC_50_ and EC_50_ were determined using a nonlinear regression.

### In vitro pMEA setup for measuring electrical spike activity (ESA)

The perforated dual MEA slice system (MEA2100; Multichannel Systems GmbH, Reutlingen, Germany) consisted of interface board (IFB-C) with integrated signal processor and headstage (HE2X60) equipped with integrated amplifiers, A/D converters and stimulus generators. A dual-perfusion system operated with the UPPER and LOWER perfusion partners running in concert to securely keep the slice adhered to the pMEA throughout the recording and to maintain a continuous inflow of fresh oxygenated aCSF ± reagent at the slice surface and a suction outflow of the exchanged bath waste through the adhered surface of the slice. Inflow was controlled by a D-Sub 9 interface, a PPS2 peristaltic perfusion pump (set at a rate of 2ml/min) fed through a cannula PH01 with integrated heating element and temperature sensor with a temperature controller interface (TC02 Multichannel Systems GmbH, Reutlingen, Germany), and a VC-8 Valve Controller (Harvard Apparatus, CT, USA). The temperature in the pMEA chamber (60pMEA200/30iR-Ti; Multichannel Systems GmbH, Reutlingen, Germany) was maintained at 32±0.25 °C for all experiments. UPPER perfusion outflow was regulated by vacuum suction through a surface-positioned metal cannula with a beveled end (Warner Instruments, ST-1 L/R 64-1401; CT, USA). LOWER inflow was driven by gravity and controlled through an IV Administration set with solution flow regulator (TrueCare Biomedix, TCBINF033G; South Miami, FL, USA). LOWER outflow was controlled by a PPS2 peristaltic perfusion pump at a rate of 0.2ml/min. Prior to starting UPPER perfusion, LOWER perfusion was adjusted in order to: (1) maintain continuous fresh aCSF/reagent exchange underneath the pMEA chamber; (2) provide gentile and constant suction to firmly contact the slice with electrodes; and (3) steadily perfuse aCSF ± reagent through the slice.

ESA detected by pMEA represented spontaneous excitatory postsynaptic field potentials (fEPSPs) and were captured simultaneously from the 60 electrodes at a sampling rate of 10 kHz within each pMEA. Data from individual electrodes were analyzed using Multi Channel Suite software (Multichannel Systems GmbH, Reutlingen, Germany). For data acquisition, Spike Detector of the MCS Experimenter was selected to Manual Threshold with defined Falling Edge of -20 mV and Rising Edge of 0 mV. Post data analysis was processed using the Spike Analyzer/MCS Analyzer. A bin width of spikes/min was obtained for spike activity analysis. Baseline recording during the first 10 min of the experiment was defined as Epoch I (0-10min), followed by Epoch II (10-20 min) through Epoch VI (50-60 min) or beyond in some cases. Since placement of each slice on pMEA was not performed with stereotactive registry among experiments, each electrode served as an independent measurement, and its temporal changes in mean spikes/min were normalized to its own baseline period (Epoch I), which served as control. Post-acquisition statistical analyses and plots of these analyses were conducted using OriginLab (OriginLab Corporation, MA, USA, 2019b). Methods used for statistical tests and normality tests are indicated in the figure/table or Supplemental Materials. Prism GraphPad (GraphPad Software, CA, USA V10) and CorelDRAW 2020 (Corel Corp.; Ottawa Canada).

### In vivo DFP dosing paradigm

Sample-size calculations based on regression analyses of preliminary data confirmed that a total of 4-5 animals per group would provide sufficient power (80+%) to detect robust effects (e.g. geometric mean ratios of 2.0+) under two-sided testing with α = 5%. Mice were injected s.c. with 9.5 mg/kg DFP in PBS (3.6 mM Na_2_HPO_4_, 1.4 mM NaH_2_PO_4_, 150 mM NaCl, pH 7.2) in the scruff of the back of the neck; vehicle controls were injected with an equal volume (0.1 ml) of PBS (s.c.). To limit peripheral parasympathomimetic toxicity, both DFP and vehicle control animals were injected 1 min later with AS (0.1 mg/kg in sterile saline, i.m.) and 2-PAM (25 mg/kg in sterile saline, i.m.) in the hind leg.

A subset of mice were randomly selected using a random number generator for treatment with one of various pharmacological antagonists of nAChR 10 min before DFP injection: the non-selective nicotinic antagonist mecamylamine hydrochloride (MEC; 9.5 mg/kg in sterile saline; AK Scientific, Union City, CA, USA), the α4-selective nAChR antagonist dihydro-β-erythroidine hydrobromide (DHβE; 5 mg/kg in sterile saline; Tocris, Bristol, UK), or the α7-selective nAChR antagonist methyllycaconitine citrate (MLA; 5 mg/kg in sterile saline; Tocris, Bristol, UK). These pharmacological reagents were injected s.c. in the scruff of the back of the neck. Vehicle control mice were injected with an equal volume (10 ml/kg) of sterile saline.

### Quantification of seizure behavior

Seizure behavior in mice was quantified using a modified Racine scale from 0 to 5 with 0 corresponding to no seizure behavior and 5 corresponding to the most severe behavior as previously described (Calsbeek et al., 2021). Seizure severity was scored every 5 min for the first 120 min after DFP injection, and then every 20 min until 240 min post-DFP. After the 4 h monitoring period, mice were weighed and then injected with 1 ml dextrose (5% v/v in saline; s.c.) in the scruff of the back of the neck and singly housed with wet food. Mice were weighed daily and provided wet food and dextrose as needed until body weight returned to pre-DFP baseline (typically within 4-7 d).

### EEG surgery and recordings

A subset of animals was instrumented for cortical electroencephalographic (EEG) monitoring in accordance with the UC Davis Rodent Survival Surgery guidelines. Adult male WT, α4 KO, and α7 KO C57BL/6J mice (8 weeks old) were deeply anesthetized using continuous isoflurane inhalation (5% for induction, 1-3% for maintenance; VetOne) in medical grade oxygen (0.5 l/min; Airgas, Radnor, PA) and compressed air (1 l/min; Airgas). Animals were secured in a stereotaxic frame (Stoelting Co., Wood Dale, IL), and the surgical site was shaved and cleaned three times with alternating betadine scrub (Purdue Products L.P., Stamford, CT) and 70% v/v isopropyl alcohol (Fisher HealthCare, Pittsburgh, PA). Sterile ophthalmic ointment (Altaire Pharmaceuticals, Northville, NY) was applied to prevent eye dryness. Analgesics (meloxicam, 2 mg/kg, s.c.) and antibiotics (ampicillin, 40 mg/kg, s.c.) were administered preoperatively.

An incision was made along the midline of the skull from the eyes to the neck, and two stainless steel head mount screws (P1 Technologies, Roanoke, VA, USA) were implanted into the skull (2 mm anterior of bregma/-1.5 mm lateral of midline; 2 mm posterior of bregma/+1.5 mm lateral of midline). Head mount screws were connected to sterile HD-X02 telemetry devices (Data Sciences International (DSI), St. Paul, MN, USA) inserted subcutaneously along the flank of the animal. The screws were further secured with a cap of dental cement, and the incision site was sutured (Ethicon Inc., Raritan, NJ). Analgesia and antibiotics were administered daily for 2 d post-surgery. Animals were allowed to recover for 7-10 d before DFP exposure. 4 of 40 mice instrumented for EEG monitoring died or were euthanized prior to experimentation.

On the day of experimentation, animals with telemetry implants were placed on PhysioTel Receivers (DSI) and baseline EEG recordings were taken for ∼ 1 h (Ponemah software, DSI) prior to DFP intoxication. EEG animals received the same DFP and nAChR antagonist dosing paradigm as non-instrumented animals, and EEG data were recorded for ∼4 h post intoxication. Animals that died during the post-intoxication monitoring period were excluded from subsequent EEG analyses.

### EEG analysis

EEG traces were analyzed using NeuroScore software, V3.3.9317-1 (DSI). Prior to subsequent analyses, traces were preprocessed using a 1 Hz high-pass filter to remove movement artifact. Researchers were blinded to experimental group during electrographic analyses. Time windows of interest were baseline (pre-DFP), early-seizure (first hour post-DFP intoxication), and late-seizure (1-4 hours post-DFP).

EEG root mean squared (RMS) value is elevated during seizure activity (Dhir et al., 2020). RMS values for EEG traces were extracted over 1-s epochs for the entire recording period. Baseline, early-, and late-seizure RMS values were determined by averaging the RMS values for each 1-s epoch over the recording window to produce a single average value for each epoch. RMS values were normalized by dividing the early- and late-seizure RMS values by baseline RMS values.

EEG spike rate is another electrographic metric consistently elevated during seizure activity (Slimen et al., 2020). Spike rate offers complementary data to RMS, capturing the frequency of abnormal coordinated discharges of neuron populations. For spike rate analysis, spike amplitude threshold criteria were determined for each animal and defined as an amplitude of greater than 4 times the standard deviation of average baseline oscillatory amplitude. For each animal, amplitude (greater than mean amplitude + 4X standard deviation) and duration criteria (20-70 ms) were used to identify individual spikes across the entire recording period. Spike count data were then binned into 1-min epochs for the entire recording period. Baseline spike rate was calculated by averaging spike count data over the recording period prior to administration of DFP. Spike rate over the early- and late-seizure periods was normalized to this baseline rate for data analysis and presentation.

Key outcomes for the statistical analysis of EEG data included RMS, normalized RMS, and spike rate. For RMS and spike rate, there were three time epochs of interest: baseline, early seizure (first hour post DFP exposure), and late seizure (1-4 hours post DFP exposure). For normalized RMS, there were two epochs chosen for analysis, early and late seizure, both relative to baseline. Due to the repeated measurements over time within an animal, mixed effects models, including animal-specific random effects, with key factors of interest including group (Veh/WT, MEC, α4 KO, α7 KO, DHβE, MLA) and time epoch were used to evaluate group differences. An interaction between the two factors was considered and Akaike Information Criterion was used to determine the best model. All outcomes were transformed using the natural logarithm to better meet the assumption of the model; spike rate values were first shifted by 0.1 prior to taking the log-transformation to account for zeros in the data. Specific contrasts for within group change (as well as differences in change between the different treatment groups and the Veh/WT group) were constructed and tested with a Wald test. The Benjamini-Hochberg false discovery rate (FDR) was used to account for multiple comparisons. Results are presented as geometric mean ratios (GMR) between time epochs within each group and between treatment groups and Veh/WT controls. Point estimates of the ratios and 95% confidence intervals are presented in the figures. When the confidence interval for the GMR includes 1, there is no statistical evidence of a difference between groups. All analyses were performed using SAS software, version 9.4 and alpha was set at 0.05; all reported results remained significant after the FDR procedure.

## Acknowledgements

We thank Dr. Suzette Smiley-Jewell (UC Davis CounterACT Center) for her assistance in editing this manuscript. We also thank Dr. Susan Hulsizer (UC Davis) for her help in setting up the multichannel system and early MEA experiments to optimize standard operating protocols of pMEA studies.

## Data Availability

For original data files, please contact, or.

## Supplemental Materials

α4/ α7 genotyping protocol details:

| Reagent | Volume (μl) |
| --- | --- |
| DNA Sample | 2 |
| Go Taq Green, Promega #M7123 | 12.5 |
| Primer 1 – stock concentration 50 μM | 0.25 |
| Primer 2 – stock concentration 50 μM | 0.25 |
| Primer 3 – stock concentration 50 μM | 0.25 |
| Primer 4 – stock concentration 50 μM | 0.25 |
| Double-distilled H<sub>2</sub>O | 9.5 |

Thermocycler Program:
| Steps | Temp ($^{\circ}$C) | Time | # of Cycles |
| --- | --- | --- | --- |
| 1. Initiation/Melting | 94 | 3 min | 1 |
| 2. Denaturation | 94 | 45 sec | 1 |
| 3. Annealing | 60 | 30 sec | 1 |
| 4. Elongation | 72 | 1 min | 1 – return to step 2 x 35 iterations |
| 5. Amplification | 72 | 10 min | 1 |
| 6. Finish | 4 | n/a | n/a |

**Supplemental Figure 1.**
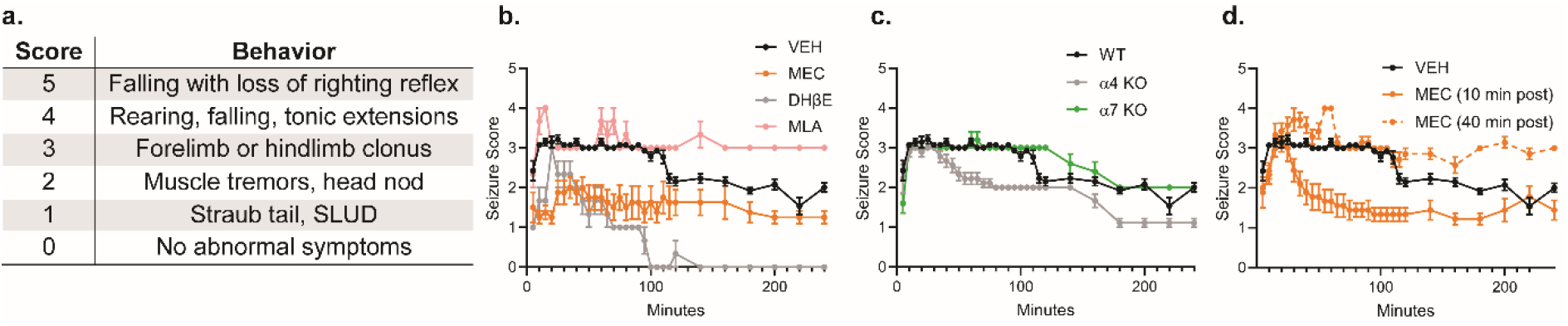
(a) Behavioral scoring scale used to evaluate seizures following DFP administration. SLUD = salivation, lacrimation, urination, defecation. (b) Average behavioral seizure scores over time following acute DFP intoxication. Mice were pretreated with vehicle (VEH), the nonselective nAChR antagonist mecamylamine (MEC), the α4-selective nAChR antagonist dihydro-β-erythroidine hydrobromide (DHβE), or the α7-selective nAChR antagonist methyllycaconitine citrate (MLA) 10 min prior to injection of DFP (9.5 mg/kg, s.c.), followed 1 min later by a combined injection of atropine sulfate (AS) (0.1 mg/kg, i.m.) and 2-pralidoxime (2-PAM) (25 mg/kg, i.m.). Data are presented as mean ± SEM (n= 3 – 14). (c) Average behavioral seizure score over time following DFP intoxication in wildtype (WT), α4 nAChR knockout (α4 KO), and α7 nAChR knockout (α7 KO) animals. Data are presented as mean ± SEM (n= 5 – 14). WT animals were the same as VEH animals presented in panel (b). (d) Average behavioral seizure score over time following DFP intoxication in VEH animals (same animals as shown in panels (b) and (c) or mice treated with MEC 10- or 40-min post-DFP intoxication. Data are presented as mean ± SEM (n= 7-14).

**Supplemental Figure 2.**
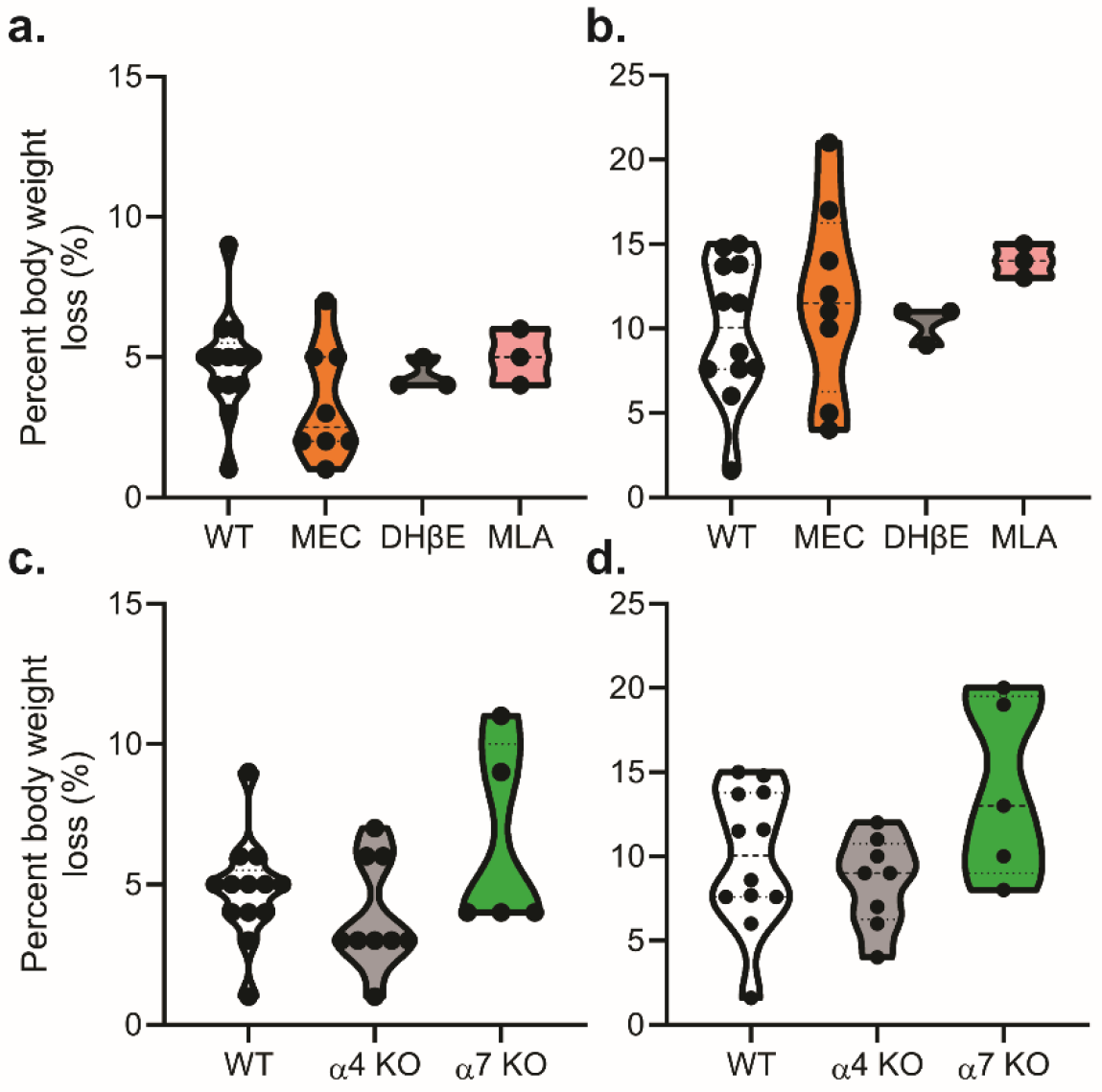
(a, b) The impact of pretreatment with nAChR antagonists on body weight loss at 4 (a) and 24 hours (b) post-DFP intoxication. (c, d) Percent body weight loss at 4 (c) and 24 hours (d) post-DFP intoxication in WT, α4 KO, and α7 KO animals. Data are presented as violin plots in which each point represents an individual animal, and the horizontal lines represent the minimum, quartile, median, and maximum values (n= 3 – 14). WT animals were the same as VEH animals presented in panels (a) and (b).

## <u>M</u>aterials <u>D</u>esign <u>A</u>nalysis <u>R</u>eporting (MDAR) Checklist for Authors

The [MDAR framework](https://osf.io/xfpn4/) establishes a minimum set of requirements in transparent reporting mainly applicable to studies in the life sciences.

*eLife* asks authors to **provide detailed information within their article** to facilitate the interpretation and replication of their work. Authors can also upload supporting materials to comply with relevant reporting guidelines for health-related research (see [EQUATOR Network](http://www.equator-network.org/)), life science research (see the [BioSharing Information Resource](http://biosharing.org/)), or animal research (see the [ARRIVE Guidelines](http://www.plosbiology.org/article/info:doi/10.1371/journal.pbio.1000412) and the [STRANGE Framework](https://doi.org/10.1038/d41586-020-01751-5); for details, see *eLife*’s [Journal Policies](https://reviewer.elifesciences.org/author-guide/journal-policies)). Where applicable, authors should refer to any relevant reporting standards materials in this form.

For all that apply, please note **where in the article** the information is provided. Please note that we also collect information about data availability and ethics in the submission form.

### Materials

**Materials:**
| **Newly created materials** | **Indicate where provided:**<br>**section/figure legend** | **N/A** |
| --- | --- | --- |
| The manuscript includes a dedicated "materials availability statement" providing transparent disclosure about availability of newly created materials including details on how materials can be accessed and describing any restrictions on access. | There are no newly created materials used in this study. | N/A |

| **Antibodies** | **Indicate where provided:**<br>**section/figure legend** | **N/A** |
| --- | --- | --- |
| For commercial reagents, provide supplier name, catalogue number and RRID, if available. |  | N/A |

| **DNA and RNA sequences** | **Indicate where provided:**<br>**section/figure legend** | **N/A** |
| --- | --- | --- |
| Short novel DNA or RNA including primers, probes: Sequences should be included or deposited in a public repository. | In Materials and Methods: Animals |  |

| **Cell materials** | **Indicate where provided:**<br>**section/figure legend** | **N/A** |
| --- | --- | --- |
| Cell lines: Provide species information, strain. Provide accession number in repository OR supplier name, catalog number, clone number, OR RRID. |  | N/A |
| Primary cultures: Provide species, strain, sex of origin, genetic modification status. | In Methods and Materials |  |

| **Experimental animals** | **Indicate where provided:**<br>**section/figure legend** | **N/A** |
| --- | --- | --- |
| Laboratory animals or Model organisms: Provide species, strain, sex, age, genetic modification status. Provide accession number in repository OR supplier name, catalog number, clone number, OR RRID. | In Methods and Materials |  |
| Animal observed in or captured from the field: Provide species, sex, and age where possible. |  | N/A |

| **Plants and microbes** | **Indicate where provided:**<br>**section/figure legend** | **N/A** |
| --- | --- | --- |
| Plants: provide species and strain, ecotype and cultivar where relevant, unique accession number if available, and source (including location for collected wild specimens). |  | N/A |
| Microbes: provide species and strain, unique accession number if available, and source. |  | N/A |

| **Human research participants** | **Indicate where provided:**<br>**section/figure legend) or state if**<br>**these demographics were not**<br>**collected** | **N/A** |
| --- | --- | --- |
| If collected and within the bounds of privacy constraints report on age, sex, gender and ethnicity for all study participants. |  | N/A |

### Design

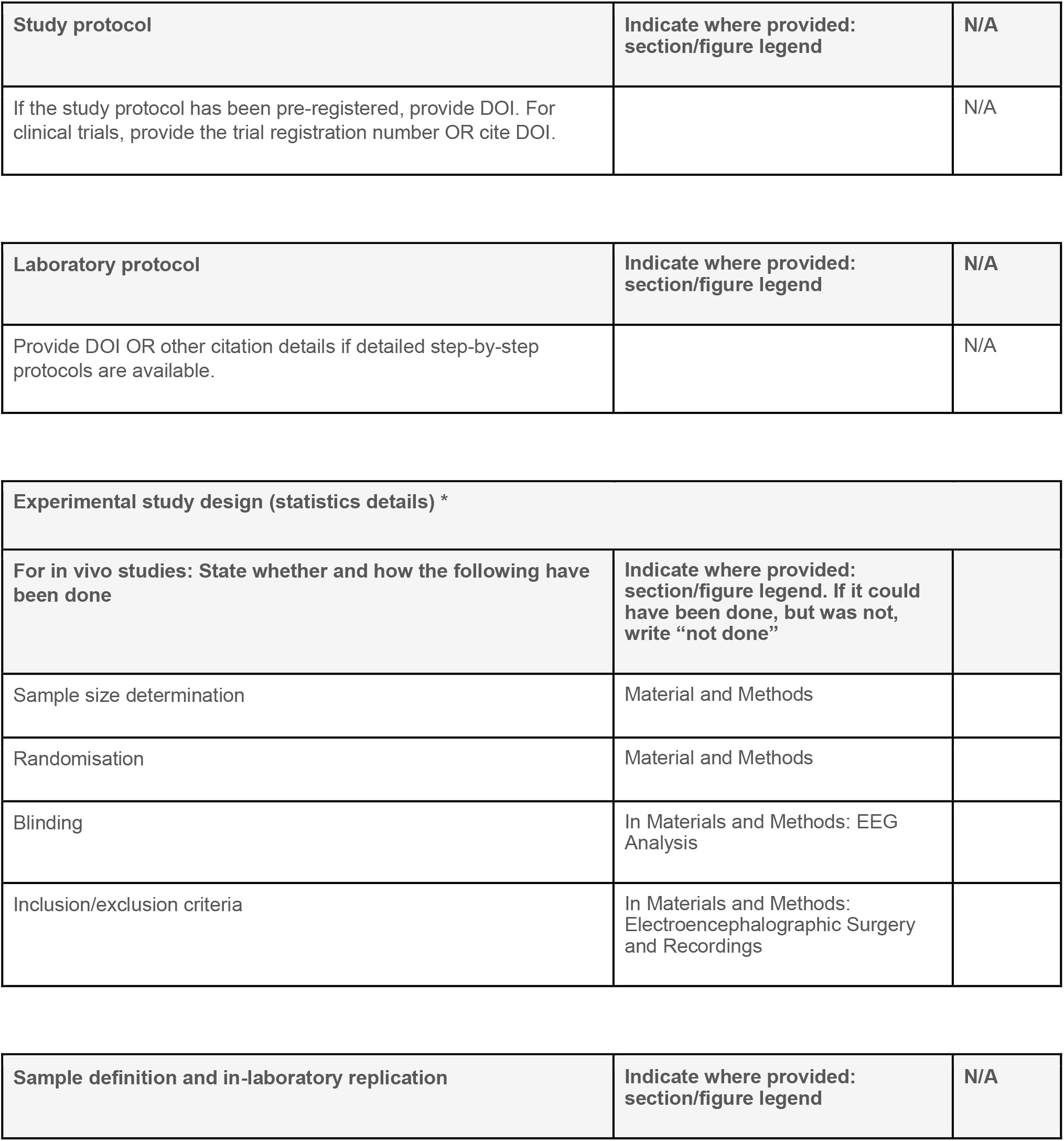
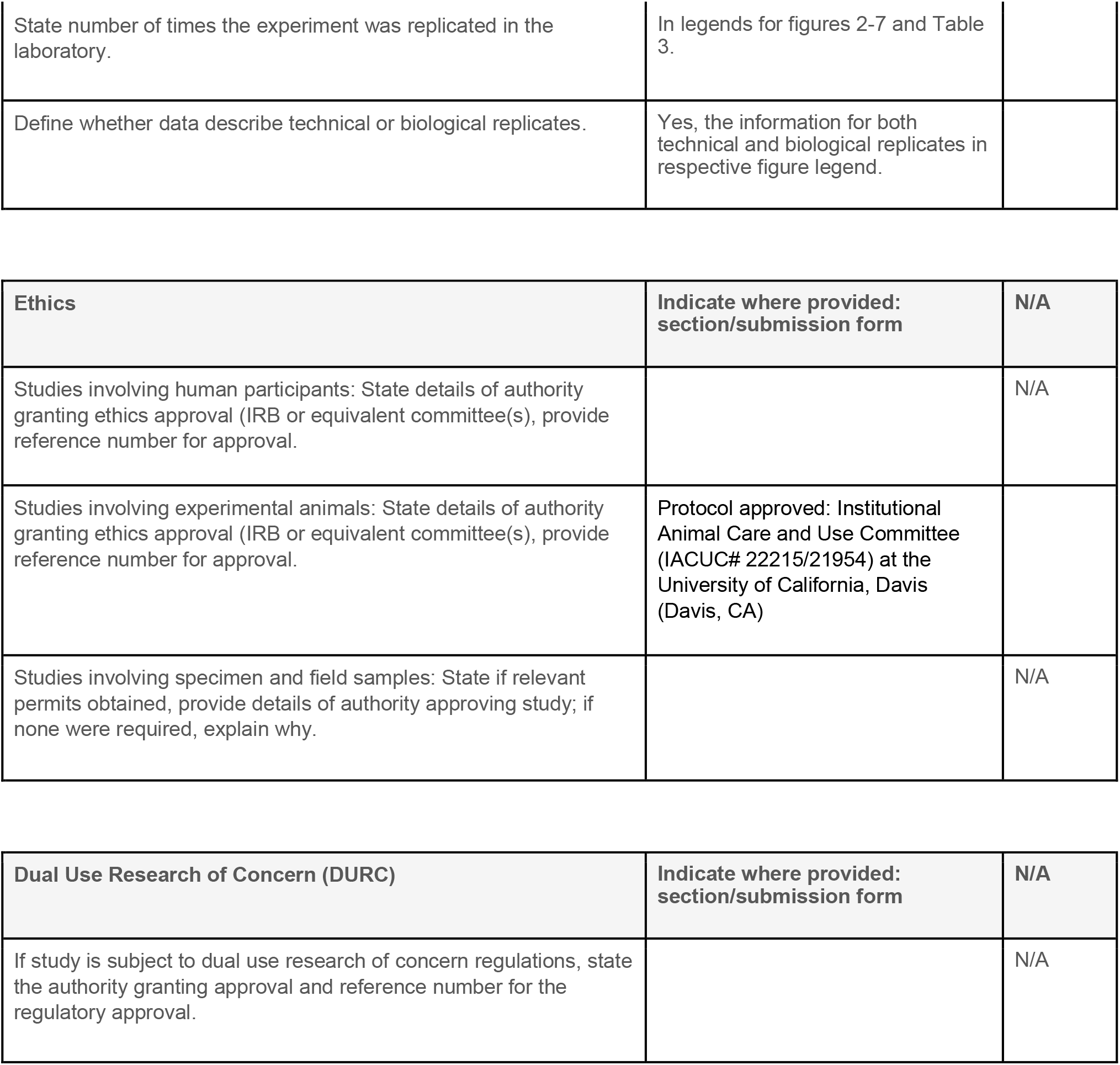

### Analysis

**Analysis:**
| **Attrition** | **Indicate where provided:**<br>**section/figure legend** | **N/A** |
| --- | --- | --- |
| Describe whether exclusion criteria were pre-established. Report if sample or data points were omitted from analysis. If yes, report if this was due to attrition or intentional exclusion and provide justification. |  | N/A |

| **Statistics** | **Indicate where provided:**<br>**section/figure legend** | **N/A** |
| --- | --- | --- |
| Describe statistical tests used and justify choice of tests. | Materials and Methods: EEG Statistical Analysis. Legends for the Table 3; figures 4-6. |  |

| **Data availability** | **Indicate where provided:**<br>**section/submission form** | **N/A** |
| --- | --- | --- |
| For newly created and reused datasets, the manuscript includes a data availability statement that provides details for access (or notes restrictions on access). | In Data Availability section |  |
| When newly created datasets are publicly available, provide accession number in repository OR DOI and licensing details where available. |  | N/A |
| If reused data is publicly available provide accession number in repository OR DOI, OR URL, OR citation. |  | N/A |

| **Code availability** | **Indicate where provided:**<br>**section/figure legend** | **N/A** |
| --- | --- | --- |
| For any computer code/software/mathematical algorithms essential for replicating the main findings of the study, whether newly generated or re-used, the manuscript includes a data availability statement that provides details for access or notes restrictions. |  | N/A |
| Where newly generated code is publicly available, provide accession number in repository, OR DOI OR URL and licensing details where available. State any restrictions on code availability or accessibility. |  | N/A |
| If reused code is publicly available provide accession number in repository OR DOI OR URL, OR citation. |  | N/A |

### Reporting

The MDAR framework recommends adoption of discipline-specific guidelines, established and endorsed through community initiatives.

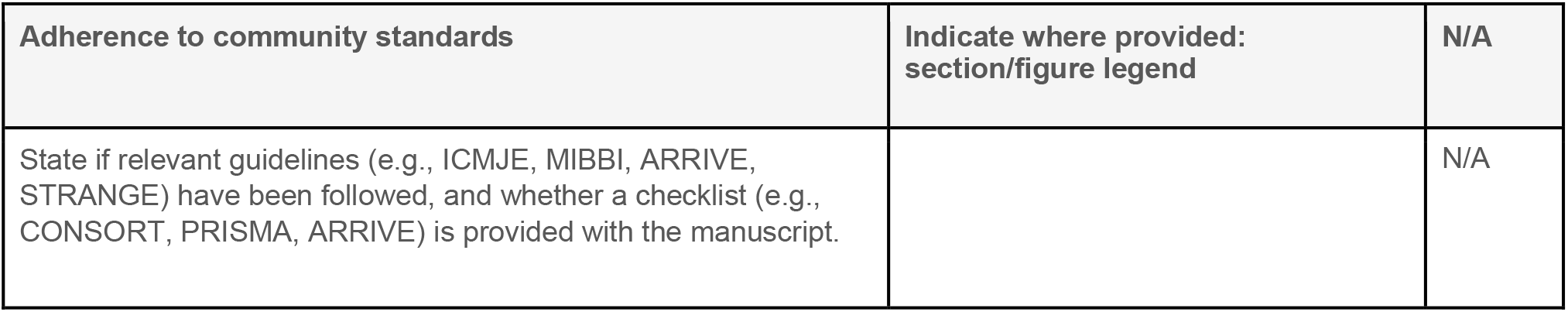

* We provide the following guidance regarding transparent reporting and statistics; we also refer authors to [Ten common statistical mistakes to watch out for when writing or reviewing a manuscript](https://doi.org/10.7554/eLife.48175).

#### Sample-size estimation

- You should state whether an appropriate sample size was computed when the study was being designed
- You should state the statistical method of sample size computation and any required assumptions
- If no explicit power analysis was used, you should describe how you decided what sample (replicate) size (number) to use

#### Replicates

- You should report how often each experiment was performed
- You should include a definition of biological versus technical replication
- The data obtained should be provided and sufficient information should be provided to indicate the number of independent biological and/or technical replicates
- If you encountered any outliers, you should describe how these were handled
- Criteria for exclusion/inclusion of data should be clearly stated
- High-throughput sequence data should be uploaded before submission, with a private link for reviewers provided (these are available from both GEO and ArrayExpress)

#### Statistical reporting

- Statistical analysis methods should be described and justified
- Raw data should be presented in figures whenever informative to do so (typically when N per group is less than 10)
- For each experiment, you should identify the statistical tests used, exact values of N, definitions of center, methods of multiple test correction, and dispersion and precision measures (e.g., mean, median, SD, SEM, confidence intervals; and, for the major substantive results, a measure of effect size (e.g., Pearson’s r, Cohen’s d)
- Report exact p-values wherever possible alongside the summary statistics and 95% confidence intervals. These should be reported for all key questions and not only when the p-value is less than 0.05.

#### Group allocation

- Indicate how samples were allocated into experimental groups (in the case of clinical studies, please specify allocation to treatment method); if randomization was used, please also state if restricted randomization was applied
- Indicate if masking was used during group allocation, data collection and/or data analysis

